# Rank-biased access to females in the likely fertile period despite comparable copulation rates in male bonobos at Wamba

**DOI:** 10.1101/2025.09.02.673851

**Authors:** Kazuya Toda, Furuichi Takeshi

## Abstract

**Objectives:** Female bonobos (*Pan paniscus*) show prolonged sexual swelling and copulate across an unusually extended timeframe, including the postpartum infertile stage. We tested the updated prolonged sexual receptivity hypothesis that the simultaneous presence of multiple receptive females disperses male mating effort and weakens male-male contest competition by examining rank effects on copulation rates and mating partners.

**Materials and Methods:** In a free-ranging bonobo group at Wamba, we collected copulation and party-composition data using all-day focal follows. Based on daily swelling scores, we operationally defined a likely fertile period (LFP) from detumescence at the end of the maximal swelling phase. We analyzed rank-dependent patterns in copulation rates and mating partners were analyzed using generalized linear mixed models (GLMMs).

**Results:** Copulation rates did not differ detectably across male dominance ranks, with approximately twice per day on average. In contrast, higher-ranking males were more likely to copulate with LFP females. At the party level, lower-ranking males increasingly copulated with non-LFP females as the number of LFP females present increased.

**Discussion:** These findings indicate that pronounced male reproductive skew in bonobos appears to arise through rank-biased access to females in the fertile window despite broadly similar copulation rates across males. Subordinate males’ tendency to shift mating effort toward non-LFP females when LFP females are available may reduce injury risks of direct contest competition while facilitating dominant males to guard fertile females effectively. Therefore, prolonged sexual receptivity in female bonobos could be a key factor yielding the condition where lower male aggression and higher reproductive skew coexist.

## 1. Introduction

Sexual receptivity can be defined as the physiological, morphological, and behavioral readiness of a mammalian female to accept a male and engage in copulation (Dixson 1998). In most non-primate mammals, receptivity is largely confined to the periovulatory phase of the estrus cycle and is tightly regulated by fluctuations in ovarian hormones such as estrogen and progesterone (Conaway 1971; Weir and Rowlands 1973). Prosimian primates show a broadly similar pattern, with female receptivity concentrated within a relatively narrow fertile window (Small 1988). In contrast, across anthropoid primates, sexual behavior is often less tightly governed by ovarian hormones, and females may remain receptive outside the fertile phase (Wallen and Zehr 2004). Prolonged sexual receptivity can impose fitness costs, including opportunity and energetic costs associated with sexual activity (Kunz et al. 2021), greater exposure to male harassment and coercion that can result in injury (Muller et al. 2011), and increased risk of sexually transmitted pathogens (Phillips-Conroy et al. 1994). Nonetheless, these costs may be outweighed when prolonged sexual receptivity benefits females, for example by confusing paternity to reduce infanticide risk and/or by eliciting paternal investment (Fernandez-Duque et al. 2020; Rooker and Gavrilets 2020).

Across anthropoids, females vary widely in whether and how they advertise fertility to males. In some polygynandrous taxa, females express graded visual cues of ovulation, such as sexual skin tumescence or skin coloration, which can bias male mating effort toward fertile periods while maintaining uncertainty across a prolonged receptive period. This combination may allow females to increase conception probability while also promoting assurance with paternity confusion (e.g., Engelhardt et al. 2004; Setchell and Wickings 2004; Brauch et al. 2007; Gesquiere et al. 2007; Dubuc et al. 2009 Higham et al. 2009). In contrast, monogamous taxa typically show slight or absent visual ovulatory cues (i.e., concealed ovulation), and females are receptive over a long range of their estrus cycles (e.g., Kendrick and Dixson 1983; Barelli et al. 2008; Fernandez-Duque et al. 2011). A comparative phylogenetic analysis suggests an evolutionary scenario that concealed ovulation among anthropoids tended to occur under polygamous or polygynandrous mating systems as a counterstrategy against male infanticide and thereafter can have facilitated the transition to monogamous (Sillén-Tullberg and Moller 1993). In humans, concealed ovulation with continuous sexual receptivity has been argued to promote pair bonds and paternal investment (Alexander and Noonan 1979; Strassmann 1981; Rodriguez-Girones and Enquist 2001). However, there are other hypotheses that concealed ovulation in humans could have evolved as a side effect of the transition to bipedal locomotion (e.g., reduced visibility of sexual swelling) or the adaptation to African savannah environment (e.g., increased physiological cost of water retention), and so on (Pawłowski 1999).

Two *Pan* species, chimpanzees (*P. troglodytes*) and bonobos (*P. paniscus*), the closest living relatives to humans, are non-seasonal breeders with interbirth intervals of approximately 4–6 years (Emery Thompson and Sabbi 2024). They live in multi-male/multi-female groups with promiscuous mating systems, where females display sexual skin tumescence as visual ovulatory cues, reflecting fluctuations in estrogen and progesterone levels (Graham et al. 1972; Heistermann et al. 1996). This advertised ovulation is phylogenetically considered to have been acquired by the common ancestor of the genus *Pan* after the divergence from the hominin ancestor (Sillén-Tullberg and Moller 1993). Although chimpanzees and bonobos diverged at an estimated 2.6–0.8 million years ago (Takemoto, Kawamoto, and Furuichi 2015; Takemoto et al. 2017), females of both *Pan* species are known to exhibit prolonged sexual receptivity from the facts that their maximal swellings persist longer than the actual fertile window in ovulation cycles and can appear after conception and during postpartum infertile phase (Deschner et al. 2003; Douglas et al. 2016).

However, female bonobos spend a remarkably longer period of displaying sexual swelling and engaging in copulation during the infertile period than female chimpanzees (Furuichi 2011). One factor reason is that the duration of maximal swelling phases in each cycle is about two days longer on average in female bonobos than chimpanzees (13.5 ± 1.8 SD days for bonobos and 11.3 ± 1.1 SD days for chimpanzees: reviewed by Ryu et al. 2025). Another factor is that the span from parturition to the first observed sexual swelling postpartum is much shorter in female bonobos than chimpanzees (0.7 ± 0.4 SD years for bonobos: Hashimoto et al. 2022; 1.8 ± 0.4 SD and 2.4 ± 1.5 SD years for western chimpanzees: Deschner and Boesch 2007; 3.4–4.6 years on average for eastern chimpanzees: Wallis 1997; Nishida et al. 2003; Emery Thompson and Ellison 2017). The physiological mechanisms underlying these interspecific differences remains poorly understood; one possibility is that the threshold of estrogenic stimulation causing sexual skin tumescence is lower in female bonobos than in female chimpanzees (Hashimoto et al. 2022).

The exceptionally early resumption of sexual swelling in female bonobos during postpartum infertile stage has been hypothesized to contribute to the increased number of receptive females in groups and thereby hinder few dominant males from monopolizing access to mating partners (termed as “prolonged sexual receptivity” hypothesis: Furuichi 2023). The operational sex ratio, calculated by the number of mature males per that of sexually receptive females present at a given time, is widely used as an indicator of the intensity of male-male contest competition over mates (Kvarnemo and Ahnesjo 1996; Weir, Grant, and Hutchings 2011). The operational sex ratio in a bonobo group at Wamba was reported to be lower than those in chimpanzee groups at Mahale and Gombe (2.8 versus 4.2 and 12.3: Furuichi and Hashimoto 2002). Moreover, although both *Pan* species exhibit a high degree of fission-fusion dynamics in their grouping patterns (Nishida 1968; Kuroda 1979), the mean number of females present within one party (i.e., a flexible subgroup whose membership varies in size and composition over time) was higher in bonobos at Wamba (3.2 ± 1.1 SD adult females) than in eastern chimpanzees at Kalinzu (1.2 ± 0.8 SD adult females) (Mulavwa et al. 2008), whereas it was only slightly higher in bonobos at LuiKotale than in western chimpanzees at Taï (4.5 versus 3.7 and 3.3 on average: Surbeck et al. 2021). High party-level association among females in bonobos may therefore further weaken male-male mating competition (Furuichi 2023).

Although male bonobos use aggression at a similar or higher frequency than male chimpanzees (Shibata and Furuichi 2024; Mouginot, Cheng, and Wilson 2023), it is widely accepted that the intensity of male aggression is substantially weaker in bonobos than in chimpanzees (Tokuyama 2023). Male chimpanzees exhibit aggressive attitudes toward males in neighboring groups and cooperate in lethal raids that can lead to territorial expansion (Watts et al. 2006; Boesch et al. 2008; Mitani, Watts, and Amsler 2010). In some populations, male chimpanzees also occasionally kill infants (Arcadi and Wrangham 1999; Wilson et al. 2014; Lowe et al. 2020) and other male within their own groups (Fawcett and Muhumuza 2000; Watts 2004; Kaburu, Inoue, and Newton-Fisher 2013). By contrast, lethal aggression has been not reported to date in male bonobos (Furuichi 2011; Wilson et al. 2014; Tokuyama 2023) but in females (Pashchevskaya et al. 2025). Skeletal evidence from museum specimens likewise indicates higher rates of trauma in chimpanzees than in bonobos (Jurmain 1997). Additionally, in 693 within-group aggressive interactions observed from a bonobo group at Wamba, there were no cases of males injuring conspecifics but two cases of female-inflicted injuries, even though males exhibit aggressive behavior more often than females (Tokuyama, Sakamaki, and Furuichi 2019). Furthermore, male bonobos have a smaller size of the canine tooth (a biological weaponry) than male chimpanzees (Kelley 1995; Balolia et al. 2025). Also, canine dimorphism, a generalized indicator of the strength of male-male contest competition over mates in primates (Plavcan and van Schaik 1992; Plavcan, van Schaik, and Kappeler 1995), is less in bonobos than chimpanzees (Kelley 1995; Balolia et al. 2025).

Previously, weak male-male mating competition in bonobos predicts that reproductive success was widely distributed among males within group. However, recent genetic paternity analyses from multiple bonobo groups at LuiKotale, Wamba, and Kokolopori consistently show pronounced male reproductive skew (Surbeck et al. 2017; Ishizuka et al. 2018; Mouginot, Cheng, and Wilson 2023). Across groups, the most successful males sired a larger share of offspring than in chimpanzees (Mouginot, Cheng, and Wilson 2023: 69 ± 11% in bonobos; 44 ± 25% in chimpanzees). Male reproductive success also covaries with dominance rank; alpha males in two study groups at Wamba achieved the highest reproductive success (Ishizuka et al. 2018), and alpha males in two of three study groups at Kokolopori sired more offspring than expected (Mouginot, Cheng, and Wilson 2023). These genetic outcomes serve as evidence that dominant males are able to monopolize access to fertile females despite noisy ovulatory signals of sexual swelling during the infertile period. Following genetic research, one recent study based on behavioral and hormonal data demonstrates that male bonobos extract usable ovulation timing with some accuracy and concentrate mating effort on females in the periovulatory phase (Ryu et al. 2025). In addition to high dominance rank empowering male bonobos to guard mates by excluding subordinate competitors (Kano 1996; Hohmann and Fruth 2003; Surbeck et al. 2012; Mouginot et al. 2024), their ability to identify fertile females by effectively managing graded ovulatory signals may help explain the pronounced male reproductive skew.

Pronounced inequality in male reproductive rates in bonobos raises the question of why male-male mating competition does not escalate into severe aggression (Shibata and Furuichi 2023). One conceivable explanation is that less aggressive males are preferred as mating partners by females or easy to receive permission of sharing the same space with with females (Surbeck et al. 2012). However, it was reported from Kokolopori bonobos that the most successful two males exhibited the highest rates of contact aggression and aggression against females more often occurred by dominant males than subordinates (Mouginot, Cheng, and Wilson 2023; Mouginot et al. 2024). This pattern suggests that female mate choice alone is unlikely to account for why severe aggression was not favored by natural selection in bonobos. Herein, we hypothesized that the simultaneous presence of multiple receptive females disperses male copulation attempts irrespective of female fertility status and thereby reduce the intensity of contest competition over mates, updating the prolonged sexual receptivity hypothesis (Furuichi 2023). Specifically. when multiple receptive females are available concurrently, subordinate males would adopt flexible mating strategies, conditionally shifting their mating effort toward females that entail lower risks of interference or aggression from dominant males. Such strategic allocation of mating effort could indirectly facilitate dominant males to monopolize access to fertile females.

It is a critical challenge to resolve the contradiction between lower male aggression and higher reproductive skew in bonobos for a better understanding of diverse social structures among primates. To examine the updated prolonged sexual receptivity hypothesis, we collected all-day focal data on copulation and party composition in a free-ranging bonobo group at Wamba. Specifically, this study addressed the following three predictions:

(1) Copulation rates are predicted to differ little among males of different ranks. If the simultaneous presence of multiple receptive females facilitates a wide range of males to engage in copulation, males of different dominance ranks would exhibit comparable copulation rates. To test this prediction, we investigated the relationship between males’ ranks and their copulation rates using all-day focal data, which reduce sampling bias relative to short focal or ad libitum sampling.
(2) Dominant males are predicted to have priority access to fertile females. Given the pronounced male reproductive skew, dominant males should copulate more frequently with fertile females than subordinates. To test this prediction, we estimated a likely fertile period (LFP) of female bonobos from the end of maximal swelling phase (MSP) based on daily assessments of sexual swelling status and compared the proportion of copulations involving higher– and lower-ranking males between LFP and non-LFP, using all-day focal data on females.
(3) Subordinate males are predicted to have a tendency to target receptive females who are less closely guarded by dominants. If males strategically allocate mating effort to reduce the risks of interference or aggression, lower-ranking males would be more likely than higher-ranking males to copulate with non-LFP females when LFP females are present in the party. To test this prediction, we analyzed male-based all-day focal data on copulation and party compositions to examine the effect of within-party male rank on the probability of copulation with non-LFP females, while distinguishing parties where LFP females were present from those where they were absent.

## 2. Materials and Methods

### 2.1 Study group and subjects

This study was conducted on a single habituated group of free-ranging bonobos (E1 group) at Wamba in the Luo Scientific Reserve, Democratic Republic of the Congo, a site studied since 1974 (Kano 1992; Furuichi et al. 2012). Data collection spanned from December 1, 2022 and May 2, 2023 (153 days). During this period, the E1 group comprised 45–48 individuals, including 10 adult males, 2 adolescent males, 17 adult females, and 3 adolescent females. The size of E1 group increased gradually after 2009, largely due to the immigration of nulliparous females (Toda, Tokuyama, and Sakamaki 2023), and thus the number of adult and adolescent females was highest ever (Figure S1). All adult and adolescent females had immigrated from other groups. Five adult and two adolescent males resided with their mothers within group.

Age classes were defined as follows. Adult males: ≥12 years, given growth in male body mass in zoo-housed bonobos ceases around 12 years (Berghaenel et al. 2023). Adolescent males: 8–<12 years, in light of the rapid rise in urinary testosterone beginning around 7 years (Behringer et al. 2014; Berghaenel et al. 2023) and the report that captive male bonobos can reproduce from about 7 years of age (Thompson-Handler 1990). Adult females: ≥12 years or parous, consistent with a mean age at first parturition of approximately 12 years at Wamba (Toda, Tokuyama, and Sakamaki 2023). Adolescent females: nulliparous females aged 8–<12 years, as females at Wamba typically disperse from the natal group at about 7 years and often engage in copulation after immigration (Toda et al. 2022).

All-day focal follows were conducted on adults and adolescents in the E1 group (Table 1: males; Table 2: females). One adult male (TN) was not followed because he had been missing from E1 for five months and was absent at the start of this study. One adult female (Aq) was also excluded from the focal subject because she had not been habituated well and thus was impossible to conduct focal follows without disturbing her natural behavior. Another adult female (No) was last seen on December 31, 2022; consequently, only one all-day focal follow was obtained for her.

**TABLE 1.** Basic information about male subjects in the E1 group (December 2022–May 2023).

| ID | Age<sup>c</sup><br>(yrs) | Resident<br>mother<sup>d</sup> | Within-sex<br>dominance rank<br>[David Score] | Focal observation<br>hours and days | Counts of copulation events during<br>focal follows |  |
| --- | --- | --- | --- | --- | --- | --- |
|  |  |  |  |  | with adults | with adolescents |
| TN<sup>a</sup> | 52* | – | Not used [-2.9] | Not collected | – | – |
| TW<sup>b</sup> | 49* | – | Not used [-12.9] | 39.2 h (4 d)<sup>e</sup> | 0 | 0 |
| DI | 48* | – | 5th [4.6] | 39.7 h (4 d) | 4 | 3 |
| NB | 35 | Ki | 4th [7.2] | 44.7 h (5 d) | 5 | 4 |
| GC | 33* | – | 8th [-11.2] | 37.5 h (4 d) | 3 | 4 |
| KT | 19 | Ki | 1st [53.4] | 36.4 h (4 d) | 3 | 0 |
| JO | 17 | Jk | 3rd [7.5] | 36.7 h (4 d) | 4 | 7 |
| KY | 14 | Ki | 2nd [10.0] | 30.6 h (4 d) | 5 | 1 |
| HC | 14 | Hs | 6th [-2.7] | 32.0 h (4 d) | 5 | 1 |
| SE | 12 | Sl | 7th [-6.1] | 32.6 h (4 d) | 7 | 2 |
| NI | 10 | Nv | 10th [-28.9] | 25.1 h (4 d) | 5 | 2 |
| YD | 9 | Yk | 9th [-27.0] | 34.4 h (4 d) | 0 | 2 |

<sup>a</sup>TN was not included in the focal subject, because he had been lost for a five month in E1 group and was absent at the beginning of this study.
<sup>b</sup>TW had seemed erectile dysfunction before the study period, and thus data on him were not used for the analysis.
<sup>c</sup>Asterisks in the column means estimated ages.
<sup>e</sup>See Table 2.
<sup>e</sup>Data in parentheses were not used for the analysis.
<sup>s</sup>Not included when female fertility status could not be determined

**TABLE 2.** Basic information about female subjects in the E1 group (December 2022–May 2023).

| ID | Age (yrs)<sup>d</sup> | Within-sex dominance rank [David Score] | Length after last parturition (months)<sup>e</sup> | Number of maximal swelling phases | Number of likely fertile periods | Focal observation hours (and days) |
| --- | --- | --- | --- | --- | --- | --- |
| No<sup>a</sup> | 52* | 12th [-2.3] | 92 | 1 | 0 | 9.0 h (1 d) |
| Ki | 49* | 1st [41.5] | 43 | 3 | 2 | 38.0 h (4 d) |
| Hs | 42* | 2nd [10.2] | 30 | 4 | 2 | 40.7 h (4 d) |
| Yk | 40* | 4th [5.3] | 16 | 1 | 0 | 35.7 h (4 d) |
| Jk | 35* | 9th [-0.29] | 28 | 4 | 3 | 38.1 h (4 d) |
| Sl | 31* | 6th [4.7] | 22 | 3 | 2 | 35.9 h (4 d) |
| Nv | 29* | 8th [2.8] | 5 | 0 | 0 | 38.3 h (4 d) |
| Ot | 26* | 10th [-0.96] | 10 | 3 | 0 | 34.5 h (4 d) |
| Fk | 25* | 5th [4.8] | 33 | 3 | 2 | 36.2 h (4 d) |
| Pf | 19* | 3rd [5.3] | 25 | 3 | 3 | 35.9 h (4 d) |
| Ik<sup>b</sup> | 16* | 15th [-6.0] | 26 | 0 | 0 | 34.5 h (4 d) |
| Sc<sup>b</sup> | 15* | 11th [-1.2] | 44 | 0 | 0 | 29.5 h (4 d) |
| Db | 14* | 7th [4.1] | 35 | 3 | 0 | 27.1 h (3 d) |
| Jj | 12* | 17th [-10.1] | 6 | 2 | 0 | 38.3 h (4 d) |
| Aq | 12* | 13th [-2.4] | 4 | 4 | 0 | Not collected |
| Ec<sup>b</sup> | 11* | 14th [-4.0] | — | 0 | 0 | 38.6 h (4 d) |
| Mz<sup>c</sup> | 10 | 18th [-10.8] | 6 | 3 | 1 | 36.5 h (4 d) |
| Tp | 10* | 20th [-11.7] | — | 5 | 0 | 35.5 h (4 d) |
| Bc | 10* | 19th [-11.3] | — | 2 | 0 | 33.0 h (4 d) |
| Ih | 9 | 16th [-8.9] | — | 4 | 0 | 34.2 h (4 d) |

<sup>a</sup>No disappeared from E1 after December 31, 2023 and is estimated to be dead due to her old age.
<sup>b</sup>Ik, Sc, and Ec were pregnant and gave birth to their infants during the study period.
<sup>c</sup>Mz had gave birth to her first infant at 9.6 years of age but lost it at 4 months after parturition.
<sup>d</sup>Asterisks in the column means estimated ages.
<sup>e</sup>At the beginning of this study period (December 6th, 2022).

### 2.2 All-day focal animal sampling

Trained research assistants followed a single party of the E1 group on a daily basis from the time the bonobos descended from their night beds (approximately 0600 h) until they constructed new night beds (about 1700 h). When the party being followed split into several parties, observers, except for those who were engaging in focal animal sampling, typically followed the largest party. The first author (KT) and a research assistant, Lokemba Batsindelia (LB), conducted focal animal sampling (Altmann 1974) throughout the day to collect continuous behavioral data and party composition on adults and adolescents.

On each observation day, we selected a focal subject from party members using a predetermined random order. A focal follow continued until one of the following conditions was met: the focal subject constructed a new night bed; darkness after sunset precluded identification; or the subject was out of sight for ≥2 consecutive hours. Focal follows with ≥4 hours of observation time were treated as all-day focal data and included in analyses. If a focal subject was lost for ≥2 consecutive hours before reaching 4 hours, a second focal follow was initiated that day only if it could start by 1200 h. If the predetermined focal subject could not be located within 30 minutes of the start of the party follow, we selected the next available individual in the predetermined order from among party members present at that time. From the midpoint of the study onward, we prioritized undersampled subjects to balance observation effort. To ensure independence among all-day focal follows, ≥20 days elapsed before resampling the same subject.

During each focal follow, we recorded a wide range of behaviors—resting, feeding, moving, and social interactions—along with the duration of each activity and the identities of participants. We also noted party composition—defined as the set of adult and adolescent individuals present during focal follows—for each time window separated at fixed hourly intervals (i.e., 0600–0700 h, 0700–0800 h, 0800–0900 h, …, 1600–1700 h), following the one-hour party method (Hashimoto, Furuichi, and Tashiro 2001). Although focal follows sometimes began before 0600 h or ended after 1700 h, 1-hour party data outside 0600–1700 h were excluded from dataset because of concern that darkness reduces the accuracy of identification.

In total, 1033.8 hours of all-day focal data were collected by KT (544.8 hours) and LB (489.0 hours) over 116 days from 11 males (384.1 hours over 45 days; Table 1) and 19 females (649.7 hours over 71 days; Table 2). Mean focal time per day was 8.9 hours (SD = 1.4; range = 4.2–11.0 hours), with a median start time of 0631 h (range = 0551–1142 h) and a median end time of 1700 h (range = 1247–1743 h). Moreover, focal party composition was recorded for a total of 1211 one-hour time windows (hereafter termed as OTWs) (males: 462 OTWs; females: 748 OTWs). Mean observation time per hourly time window was 51.8 minutes (SD = 13.3, range = 0.2–60.0 minutes).

### 2.3 Definition of copulation

Copulation was defined as a heterosexual interaction in which a male mounted a female with penile-vaginal intromission accompanied by thrusting. Because ejaculation was often not visually discernible, intercourse was recorded as copulation regardless of whether ejaculation could be confirmed (Hashimoto 1997; Ryu et al. 2025). We recorded all observable copulatory interactions between adolescent or adult individuals with ad libitum sampling. To avoid inflating counts, successive copulations involving the same dyad within 10 minutes were treated as a single copulation event.

During all-day focal follows, we observed 67 copulation events involving the focal male and 67 involving the focal female. In addition, we recorded 134 copulations ad libitum that did not involve the focal subject, yielding 268 copulations in total (19–38 events per male). Visible sperm expulsion from the vagina immediately after copulation was confirmed in 17/268 events (6.3%). With the exception of one older male (TW; estimated 51 years old), copulation was observed from all male subjects. TW has not been observed copulating or exhibiting erections since at least 2015 (Toda personal observation), and thus his focal data (38.5 hours and 42 OTWs) were excluded from analyses.

### 2.4 Definition of aggression

Aggression was defined as a directed aggressive interaction, including contact aggression (e.g., punching, kicking, pushing, tackling, biting) and threat display (e.g., chasing, charging, charging while branch-dragging, branch-shaking) (Nishida et al. 1999). Undirected display lacking a clear target was not classified as aggression. We recorded all aggressive interactions between adolescent or adult individuals and noted the target’s response, including submissive behavior (e.g., fleeing, grinning, screaming) and retaliation toward the aggressor. Successive aggressive interactions involving the same dyad within 10 minutes were treated as a single aggression event.

In total, we recorded 297 aggression events: 124 involving the focal male, 51 involving the focal female, and 122 not involving the the focal subject. Of these, 138 were male-male (46.5%), 81 male-to-female (27.3%), 62 female-to-male (20.9%), and 16 female-female (5.4%). Contact aggression occurred in 95/297 events (32.0%): initiated by males in 43/95 (45.2%) and by females in 52/95 (54.7%). Submissive responses were observed in 201/297 events (67.7%), allowing us to identify winners and losers.

### 2.5 Evaluation of male dominance ranks

To infer the dominance hierarchy of the E1 group, we calculated David’s scores (David 1987) from an interaction matrix comprising 201 aggressive interactions that elicited submissive responses, using dyadic dominance indices that incorporate win-loss proportions and correct for unequal interaction opportunities (de Vries 1998). Although our primary aim was to evaluate male dominance ranks, both sexes were included in the same matrix because male bonobos are codominant with females (Furuichi 1997; Surbeck and Hohmann 2013). We computed the David’s scores for each male (*N* = 12; Table 1) and each female (*N* = 20; Table 2) with the ‘DS’ function in the ‘EloRating’ package (Neumann and Kulik 2024) in R version 4.4.2 (R Core Team 2024). We also quantified hierarchy steepness based on David’s scores (de Vries, Stevens, and Vervaecke 2006) and the modified Landau’s linearity index (de Vries 1995), and compared the observed values with those from randomized matrices using the ‘steepness’ and ‘h.index’ functions in the ‘EloRating’ package. Both steepness and linearity were greater than expected (one-tailed *p* = 0.001 and *p* = 0.020, respectively).

The median rates of aggression from the focal male during all-day focal follows were 0.067 events/h towards males (range = 0–0.332, *N* = 32) and 0.052 events/h towards and females (range = 0–0.221, *N* = 31). The former figure is quite similar as the frequency reported from Mouginot et al. (2024) at Kokolopori (median = 0.08 events/h, range = 0–0.300) but the latter figure appears about 10 times higher (median = 0.005 events/h, range = 0–0.027), yet aggression elicited submissive responses from females was in the minority (11/31 events [35.5%]). The correlation of male dominance ranks with the rate of aggression towards males was not significant (Spearman’s rank correlation test: ρ = –0.55, *p* = 0.100), but that towards females was significant (ρ = –0.74, *p* = 0.015).

### 2.6 Assessment of sexual swelling states

We recorded the daily variation in sexual swelling states of female bonobos in the E1 group, by scoring the size and firmness of the sexual skin. Following established protocols (Furuichi 1987; Ryu, Hill, and Furuichi 2015), each female was assigned a score of 1 to 3: 1 = no swelling, 2 = intermediate swelling, 3 = maximal swelling (Ryu et al. 2025). Score 1 was determined with the minimum range of sexual skin size. However, the size alone did not always clearly separate scores 2 and 3, we additionally relied on the firmness and shininess seen of the sexual skin to distinguish these categories. Detumescence day was defined as the first day on which the maximal swelling began to subside, accompanied by loss of firmness and shininess, and the score shifted from 3 to 2.

To reduce observation error and scoring bias, researchers and trained assistants discussed and reached consensus on each female’s swelling score at a daily evening meeting. During the study period, the mean number of females with maximal swelling in the followed party per observation day was 4.5 (SD = 2.4, range = 0–10), comprising 3.1 adult females (SD = 1.8, range = 0–7, *N* = 16) and 1.4 adolescent females (SD = 1.0, range = 0–3, *N* = 3).

The maximal swelling phase (MSP) was defined as the interval from the first day a female was scored 3 to the last day scored 3 immediately before detumescence. Sample gaps in daily scores arose when a female was absent from the followed party or when observations were missed. If the score was identical immediately before and after a sample gap of ≤ 3 days, we interpolated the same score across the gap. If the score dropped from 3 to 2 and returned to 3 within 4 days, the intervening period at score 2 was considered part of a continuous MSP, following previous studies (Douglas et al. 2016; Ryu et al. 2025). Over the study period, we identified 48 MSPs from 13 adult females and 3 adolescent females (Table 2).

### 2.7 Estimation of female fertile windows

The relationship between ovulation timing and the maximal swelling phase (MSP) in female bonobos has been examined using ovarian steroid profiles measured from urine or fecal samples. Some studies suggest that ovulation can occur outside the MSP and that its timing is difficult to predict from MSP onset or end (Reichert et al. 2002; Douglas et al. 2016). In contrast, other work indicates that MSP end tends to occur a few days after ovulation and that this temporal relationship is less variable (Heistermann et al. 1996; Ryu et al. 2025). In field studies of non-human primates, the fertile phase is often operationally defined as a 4-day window comprising the day of ovulation and the three preceding days (Deschner et al. 2003; Douglas et al. 2016). This definition is biologically motivated by the short post-ovulatory viability of the ovum (generally < 24 h) and evidence from human fertility studies indicating that most conceptions result from intercourse occurring within a few days prior to ovulation, with comparatively fer pregnancies attributable to sperm older than three days (Wilcox, Weinberg, and Baird 1995; Dunson et al. 1999). Applying this criterion, Ryu et al. (2025) suggested that the probability of sperm-ovum overlap (i.e. fertility) remains high from 7 to 2 days before detumescence (peaking 4–5 days before detumescence) and then drops to zero by the day before detumescence. Ryu et al. (2025) also demonstrated that male bonobos concentrated mating effort during the late MSPs and largely ceased such effort after detumescence.

It was impractical to implement the intensive sampling necessary to pinpoint ovulation for each female in parallel with full-day focal follows. Instead, we operationally defined the likely fertile period (LFP) as the 6-day window from 7 to 2 days before MSP end in ovulatory cycles, following Ryu et al. (2025), which was conducted at the same field site and using the same visual assessment method of sexual swelling states. Nonetheless, MSPs do not necessarily coincide with ovulation (Douglas et al. 2016; Ryu et al. 2025), and distinguishing from ovulatory from anovulatory cycles based on visual records of swelling states alone is difficult. To increase the validity of the LFP assignment, we applied the following criteria.

[1] Early adolescent females were excluded from LFP candidates. In chimpanzees, adolescent females experience a prolong period of subfecund swelling cycles after menarche, likely due to anovulatory cycles and early abortions associated with an immature hypothalamic-pituitary-ovarian axis (Young and Yerkes 1943; Wallis 1997; Nishida et al. 2003). Captive bonobos also reportedly reach menarche at 8.1 ± 0.6 years (*N* = 3) and experience 4.4 ± 2.6 years of adolescent subfecundity (*N* = 5) (Thompson-Handler 1990). Accordingly, we classified 11 MSPs from 3 adolescent females (Tp, Bc, Ih) younger than 11 years into anovulatory.
[2] Adult females in early lactation were excluded from LFP candidates. In the E1 group, the mean interbirth interval is 57.7 ± 9.6 months (range = 43–72, *N* = 23) (Hashimoto et al. 2022). However, recently, the shorter interbirth interval at 31 and 32 months has been recorded at Wamba (unpublished data). Subtracting gestation length (7.9 months; Furuichi et al. 1998) suggests her conception at approximately 23–24 months postpartum. In addition, the earliest ovulation estimated from urinary hormone profiles was confirmed to be 24 months after parturition from a female in E1 (Hashimoto et al. 2022). Based on these early resumption cases, we classified 6 MSPs from 5 adult females (Yk, Nv, Ot, Jj, Aq) with infants younger 24 months into anovulatory. On the other hand, ovulation may resume rapidly after infant loss: one female that lost her infant gave birth 9 months later (Cheng et al. 2022), implying potential resumption within about a month of infant death. This study included one female (Mz) losing her infant
[3] MSPs of short duration were considered as anovulatory. At Wamba, MSP duration in ovulatory cycles (mean = 20.7 ± 6.4 SD, range = 14–38, *N* = 14) appears longer than in all cycles combined (mean = 14.0 ± 11.2 SD, range = 1–45, *N* = 36) (Ryu 2017). We therefore classified 5 MSPs lasting <8 days (mean ovulatory MSP duration minus 2 SD, based on Ryu 2017) into anovulatory.
[4] MSPs followed by an unusually short interval to the next MSP were considered as anovulatory. In female bonobos, luteal-phase length, which reflected by progesterone from the corpus luteum and associated detumescence, lasts approximately 9–13 days (Heistermann et al. 1996: 13.0 ± 0.7 SD, *N* = 6; Douglas et al. 2016: 9.5 ± 1.2 SD, *N* = 6; Ryu 2017: 10.6 ± 0.6 SD, *N* = 3). Accordingly, we classified 2 MSPs with an interval of <9 days between MSP and and the onset of the next MSP as of interval to next MSP onset (mean luteal phase length minus 2 SD, based on Ryu 2017) into anovulatory.

Overall, we classified 20/48 MSPs as ovulatory cycles in 9 adult females (No, Ki, Hs, Jk, Sl, Fk, Pf, Db, Mz) (Table 2). However, in 5 MSPs, the exact day of detumescence could not be determined because it fell within a sample gap. When detumescence was obscured by a gap, females were treated as having indeterminate LFP status during the 6 days of the MSP preceding the gap, whereas earlier days in that MSP were classified as non-LFP. Copulation with adult or adolescent males was observed at least once during 14/15 LFPs.

### 2.8 Data analyses

Statistical analyses were conducted in R version 4.4.2 (R Core Team 2024) using RStudio version 12.0.467 (Posit Team 2024). We fitted generalized linear mixed models (GLMMs) to account for repeated measures within individuals (Bolker et al. 2009) with the ‘glmer’ function in the ‘lme4’ package (Bates et al. 2015). All continuous predictors were z-transformed (mean = 0, SD = 1) to improve the model convergence, performance, and interpretability (Schielzeth 2010). Variables included in each model are described below.

To test the first prediction—that male copulation rates differ little across dominance ranks—we examined the effect of male dominance ranks on copulation rates. This analysis used all-day focal data on males (345.6 hours across 41 days), excluding one old male (TW). Because adult male chimpanzees, particularly high-ranking males, show lower sexual interest in adolescent than adult females (Muller, Thompson, and Wrangham 2006; Reddy et al. 2021), we tallied, for each focal day, the focal male’s copulations with adult and adolescent females separately. These daily counts were analyzed using two Poisson GLMMs using a log link, each including an offset for observation time to account for variation in focal follow duration (Zuur et al. 2009). Male rank (1–10) was included as a predictor of interest. In addition, the number of non-mother females with maximal swellings present on each focal day (adults: 0–6; adolescents: 0–4) was included as a control predictor to capture day-to-day variation in copulation opportunities. A random intercept for focal male identity was included in both models.

To test the second prediction—that higher-ranking males copulate more frequently with LFP females than lower-ranking males—we examined the effect of female LFP status on the probability of copulation with higher-versus lower-ranking males. All-day focal data on females included 22 days (194.8 hours) during which at least one copulation event was observed. After excluding 2 days (17.6 hours) in which the focal female’s LFP status was indeterminate, we aggregated for each remaining day (*N* = 20 days), counts of copulations with high-ranking males (ranks 1–5) and low-ranking males (ranks 6–10). These paired counts were fitted as a two-column response using a binomial GLMM with a logit link (Zuur et al. 2009). Female LFP status (LFP: *N* = 4 days; non-LFP: *N* = 16) was included as a predictor of interest, with a random intercept for focal female identity. Also, the ratio of numbers of high-versus low-ranking males observed during female focal follows was also included as a control predictor.

To reinforce the result of the analysis above at which males were roughly divided into two rank classes (higher and lower), we additionally examined the effect of male dominance ranks on the proportion of copulations with LFP versus non-LFP females using all copulation events (*N* = 268) recorded through both focal and ad libitum sampling. Of them, we excluded 11 copulations involving females of indeterminate LFP status, we aggregated, for each male, counts of copulations with LFP females (*N* = 59 events) and non-LFP females (*N* = 198 events). These two-column responses (LFP, non-LFP) were analyzed using a binomial GLM with a logit link (Zuur et al. 2009), with male rank (1–10) included as a predictor of interest.

To test the third prediction—that subordinate males selectively target females who are less closely guarded by dominant males as mating partners—we assessed the effect of within-party male dominance ranks on the probability of copulation within each one-hour time window (OTW) during male focal follows. Specifically, we fitted three separate binomial GLMMs to examine (i) whether higher-ranking males within the focal party were more likely to copulate, (ii) whether higher-ranking males within the focal party were more likely to copulate with LFP females when LFP females were present, and (iii) whether lower-ranking males within the focal party were more likely to copulate with non-LFP females when LFP females were present. Herein, we excluded OTWs in which the focal male was out of sight for >30 minutes (*N* = 41), because the number of bonobos was reported to increase with the amount of observation time until 30 minutes and then become saturated (Mulavwa et al. 2008). Also, OTWs in which the focal male was resting for >95% of observation time (*N* = 19) were excluded to eliminate datapoints where the focal male was almost inactive by lying on the bed. For each OTW, copulation occurrence was coded as 1 (≥1 copulation involving the focal male) or 0 (on copulation) and analyzed using binomial GLMMs. Copulation spanning consecutive OTWs were assigned to the earlier OTW based on the start time of the interaction. A random intercept for focal male identity was included in all models. Datasets and predictors for each model were as follows.

(i). The first model used 360 OTWs collected over 41 days, including 58 OTWs with copulation and 302 without. The number of males higher-ranking than the focal male within the party (0–7), representing the focal male’s within-party rank position, was included as the predictor of interest. The numbers of non-mother non-LFP females with maximal swelling (0–7) and non-mother LFP females (0–2) in the party were included as control predictors.
(ii). The second model used 117 OTWs collected over 17 days in which LFP females were present, including 10 OTWs with copulation with LFP females and 107 without, excluding 31 OTWs in which females of indeterminate LFP status were present. Predictors included the numbers of males higher-ranking than the focal male (0–7), non-mother non-LFP females with maximal swelling (0–7), and non-mother LFP females (1–2) within the party.
(iii). The third model used 329 OTWs collected over 38 days, where the LFP status of all females were determined, including 42 OTWs with copulation with non-LFP females and 287 without. In addition to the numbers of males higher-ranking than the focal male (0–7), non-mother non-LFP females with maximal swelling (0–7), and non-mother LFP females (0–2), we included an interaction between the number of higher-ranking males and the number of non-mother LFP females.

For model selection, we compared the goodness of fit of a single candidate model containing predictors of interest with that of the corresponding null model retaining the control predictor and random-effects structure (Bolker et al. 2009), following a null-hypothesis significance testing framework (Nickerson 2000). Model comparisons were performed using simulated likelihood ratio tests based on parametric bootstrap (1000 iterations) implemented with the ‘simulateLRT’ function in the ‘DHARMa’ package (Hartig 2024). Fixed-effects estimates were interpreted only when the candidate model significantly outperformed the null model. Confidence intervals for fixed effects were obtained via parametric bootstrap inference (2000 iterations) using the ‘bootMer’ function in the ‘lme4’ package. For each fitted model, residual diagnostics, including normality, dispersion, and the presence of outliers, were evaluated (Harrison et al. 2018) using the ‘simulateResiduals’ function in the ‘DHARMa’ package (Hartig 2024); no violations were detected (Figures S2–S8). Residual independence was assessed using the ‘check_autocorrelation’ function in the ‘performance’ package (Lüdecke et al. 2021); no significant autocorrelation was detected (minimum *p* = 0.218). Multicollinearity was evaluated with the ‘check_collinearity’ function in the same package (Lüdecke et al. 2021); variance inflation factors (VIFs) were low in all models (maximum VIF = 2.11).

### 2.9 Ethical statement

The field studies of wild bonobos were approved by the Ministry of Scientific Research and the Center for Research on Ecology and Forestry in the Democratic Republic of Congo. This study conformed to the Animal Research Guidelines in The Graduate University for Advanced Studies, Sokendai, Japan, revised on March 23, 2021.

## 3. Results

### 3.1 Relationship between male copulation rate and dominance rank

On the majority of male focal days (38 of 41 days, 92.7%), at least one non-mother female in the party exhibited maximal swelling. In addition, at least one non-mother female in the likely fertile period (LFP) was present in 17 male focal days (41%). The mean number of non-mother females with maximal swellings per day was 4.1 ± 2.4 SD (2.4 ± 1.9 SD for adult females and 1.6 ± 1.2 SD for adolescent females, respectively). Copulations were observed on 33 of 38 days (86.8%) at which maxima swelling females are present. Across males (*N* = 10), the mean copulation rate was 0.20 ± 0.08 events per focal observation hour (range = 0.06–0.29, *N* = 10 males; Table 1), comprising 0.12 ± 0.06 events/h with adult females (range = 0–0.21) and 0.07 ± 0.05 events/h with adolescent females (range = 0–0.19). Of 83 male-male aggressive interactions involving the focal male, only 5 cases (6.0%) occurred in mating contexts when targets approached or solicited maximally tumescent females or immediately after copulation.

Poisson GLMMs testing the effect of male rank on copulation counts did not differ from their respective null models for either adult females (log-likelihood ratio = 0.19, *p* = 0.503) or adolescent females (log-likelihood ratio = 0.13, *p* = 0.623). These results indicate that male copulation rates were not dependent on dominance rank, consistent with the first prediction that male copulation rates are broadly comparable across ranks.

### 3.2 Male dominance rank and access to females in the likely fertile period (LFP)

Mating partners differed markedly according to the LFP status during female focal follows. Among 15 copulations involving two LFP females, 13 events (86.7%) involved high-ranking males (ranks 1–5), and 5 events (33.3%) involved the alpha male. In contrast, among 48 copulations involving nine non-LFP females, 28 events (41.7%) were with high-ranking males, and only 2 events (4.2%) involved the alpha male.

A binomial GLMM examining the effect of female LFP status on the proportion of copulations with high-versus low-ranking males significantly outperformed the null model (log-likelihood ratio = 3.3, *p* = 0.014). Females in the LFP were therefore more likely to copulate with high-ranking males than were non-LFP females (Table 3A; Figure 1), supporting the second prediction that high-ranking males have priority access to LFP females.

**Figure 1.**
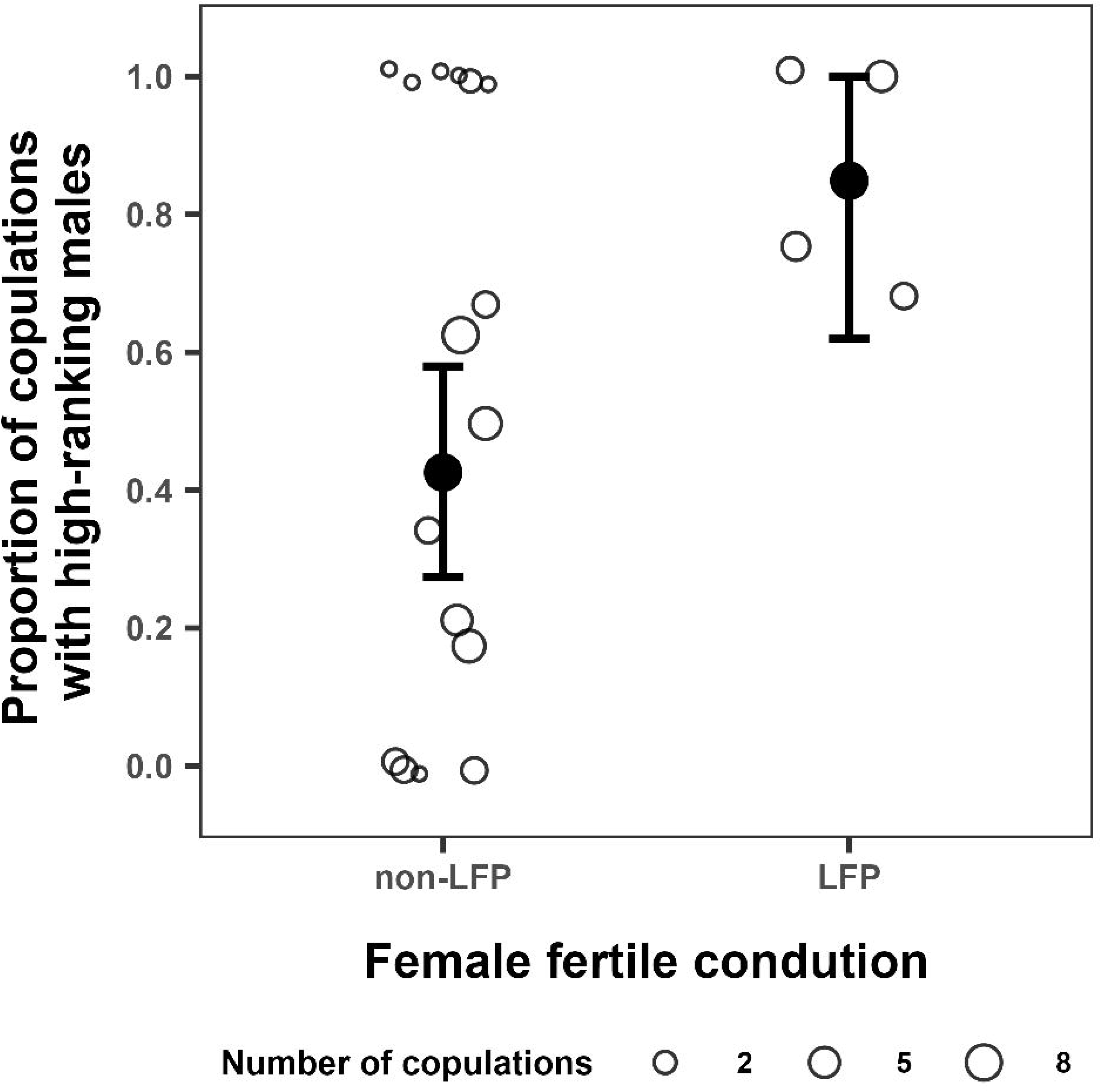
Comparisons in the proportion of copulations involving high-ranking males between females in the likely fertile period (LFP) and non-LFP females. Open circles represent observed proportions for each female focal follow, with circle size indicating the total number of copulations. Filled circles denote model-estimated means, and vertical error bars indicate confidence intervals.

**TABLE 3.**
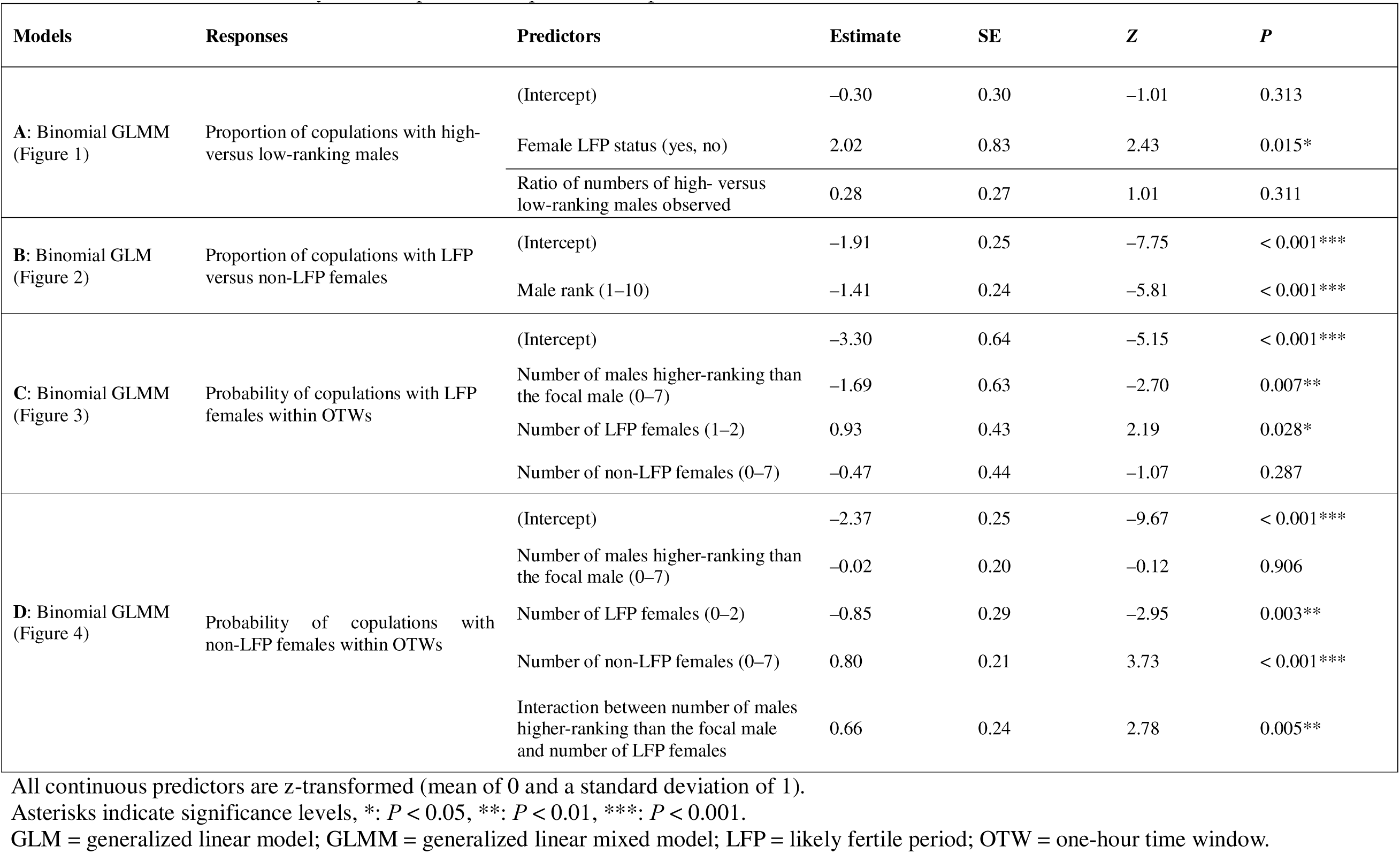
Results of model analyses for copulation frequencies and patterns.

An additional analysis of all copulation events for which female LFP status could be estimated (*N* = 257) further supported this copulation pattern. A binomial GLM examining the effect of male rank on the proportion of copulations with LFP versus non-LFP females significantly outperformed the null model (log-likelihood ratio = 25.9, *p* < 0.001). Higher-ranking males were more likely than lower-ranking males to copulate with LFP females (Table 3B; Figure 2).

**Figure 2.**
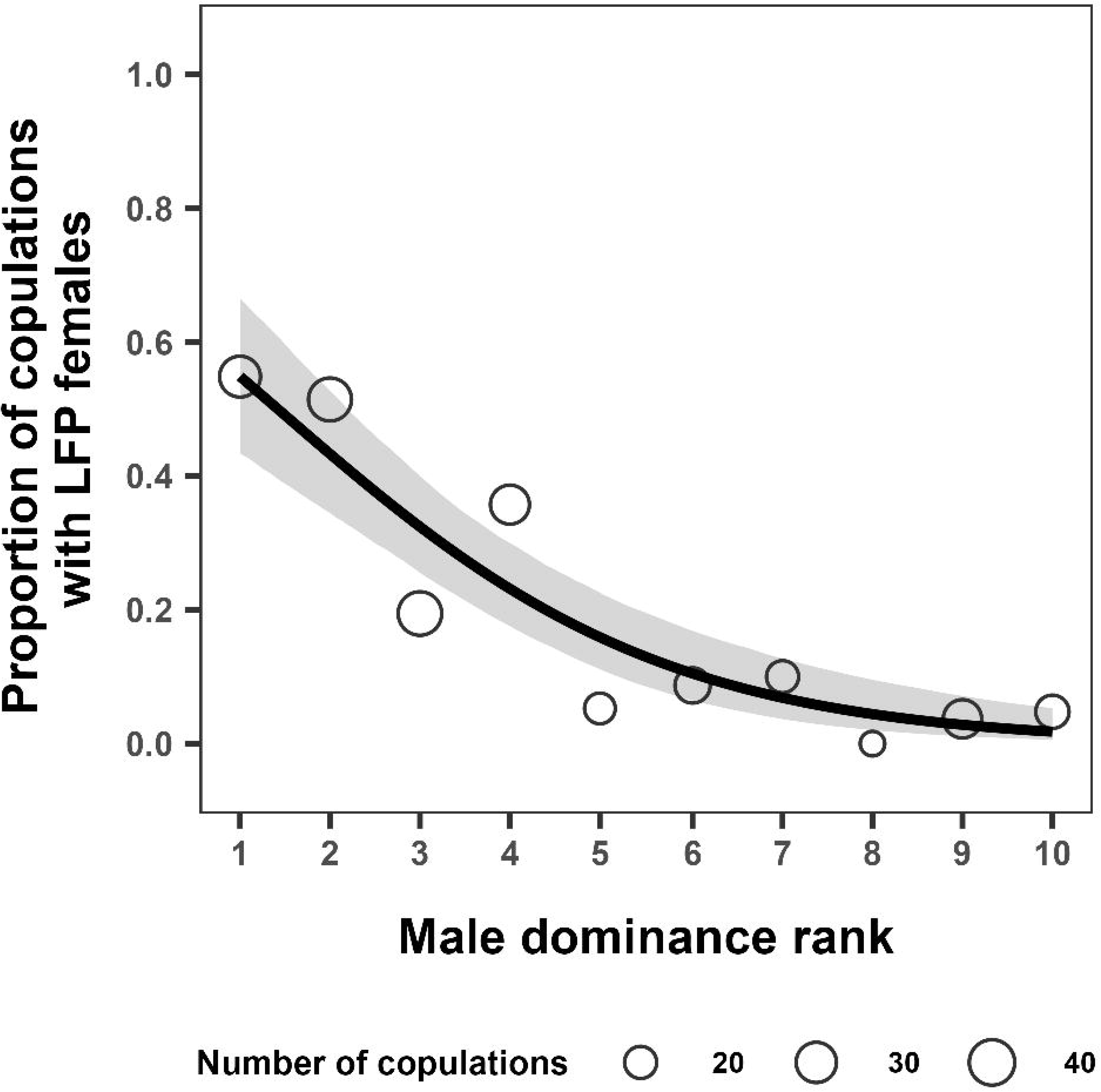
Relationship between male dominance rank and proportion of copulations involving females in the likely fertile period (LFP). Male dominance rank is ordered such that lower values indicate higher-ranking males and higher values indicate lower-ranking males. Open circles represent observed proportions for each male, with circle size indicating the total number of copulations. The solid line shows the model-predicted relationship, and the shaded area represents the 95% confidence interval.

As for all copulation events, the median proportion of copulations with LFP females across males was 9.3% (range = 0–54.8%, N = 10 males). The alpha male (KT) copulated with LFP females in 54.8% of his copulations. His maternal brothers (KY and NB), sons of the alpha female (Ki) (Table 2), also showed higher proportions (51.4% and 35.7%, respectively) than the across-male median. We observed Ki supported her sons in aggressive interactions with other individuals than maternal brothers on four occasions; including one instance in which she intercepted another male’s copulation attempt. The dyad most frequently involved in aggressive interactions was the maternal brothers, KT and KY (27 of 297 events [9.1%]), and all of them were directed from the older (KT) to younger (KY) brothers. We also observed Ki intervened in their aggressive interactions on six occasions and seemingly tried to stop the aggression of KT.

Among males whose mothers were not top-ranking, the third-ranking male (JO) copulated with LFP females at a higher proportion (19.4%) than did four lower-ranking males with resident mothers (HC, SE, YD, and NI; range = 3.7–10.0%). However, the mother of JO (Jk) did not support her son in aggressive interactions with other males throughout the study period. Two males without resident mothers (DI and GC) copulated with LFP females once or not at all.

Among all copulation events, sperm expulsion was confirmed in 6 of 59 copulations with LFP females (10.2%) and in 8 of 162 with non-LFP females (4.9%). Ejaculation was therefore observed approximately twice as frequently during copulations with LFP females as during copulations with non-LFP females, although this difference was not statistically significant (Fisher’s Exact Test: *p* = 0.209).

### 3.3 Effects of within-party male dominance ranks on mating partners

One-hour time windows (OTW) during male focal follows (*N* = 391) consisted of 4.4 ± 2.4 males (range = 1–9) and 9.1 ± 4.1 females (0–19). The mean number of non-mother females with maximal swellings was 2.9 ± 1.9 females (range = 0–9), consisting of 2.5 ± 1.7 non-LFP and 0.4 ± 0.6 LFP females. Of these OTWs, at least one non-LFP female with maximal swelling was present in 354 OTWs (90.5%), and at least one LFP female was present in 129 OTWs (33.0%).

The first model examining the effect of the number of males higher ranking than the focal male (i.e., within-party male rank) on the occurrence of copulations within one-hour time windows (OTWs) did not outperform the null model (log-likelihood ratio = 1.6, *p* = 0.075). This result indicates that within-party male rank alone did not affect the probability of copulation, following the first prediction.

In contrast, the second model examining the effect of within-party male rank, expressed as the number of males ranking higher than the focal male within the party, on the occurrence of copulations with LFP females within OTWs significantly outperformed the null model (log-likelihood ratio = 5.7, *p* = 0.001). Within-party higher-ranking males were more likely than lower-ranking males to copulate with LFP females when such females were present in the party (Table 3C; Figure 3), consistent with the second prediction.

**Figure 3.**
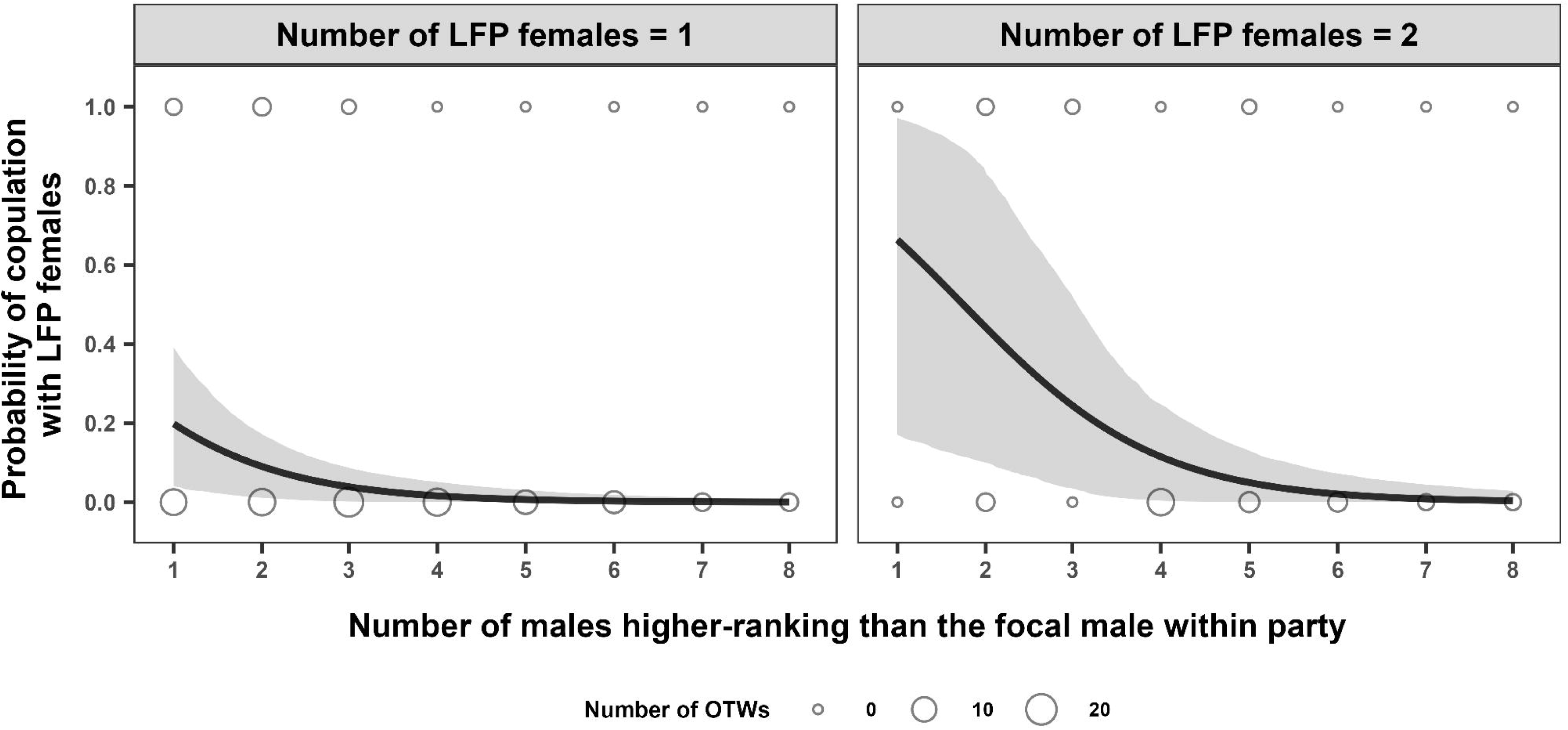
Relationship between within-party male dominance rank and the probability of copulation with females in the likely fertile period (LFP). Predicted copulation probabilities are shown separately for one-hour time windows (OTWs) containing one (left panel) or two (right panel) females in the LFP. The size of open circles indicates observed total counts of whether copulation occurred for each OTW. Solid lines show model-predicted probabilities, and shaded areas represent 95% confidence intervals on the probability scale.

Finally, the third model examining the interaction between within-party male rank and the number of LFP females present on the occurrence of copulations with non-LFP females within OTWs significantly outperformed the null model (log-likelihood ratio = 7.3, *p* = 0.002). The significant interaction indicates that, as the number of LFP females increased, within-party lower-ranking males were more likely than higher-ranking males to copulate with non-LFP females (Table 3D; Figure 4). This pattern supports the third prediction that subordinate males selectively target non-LFP females as mating partners when LFP females are available.

**Figure 4.**
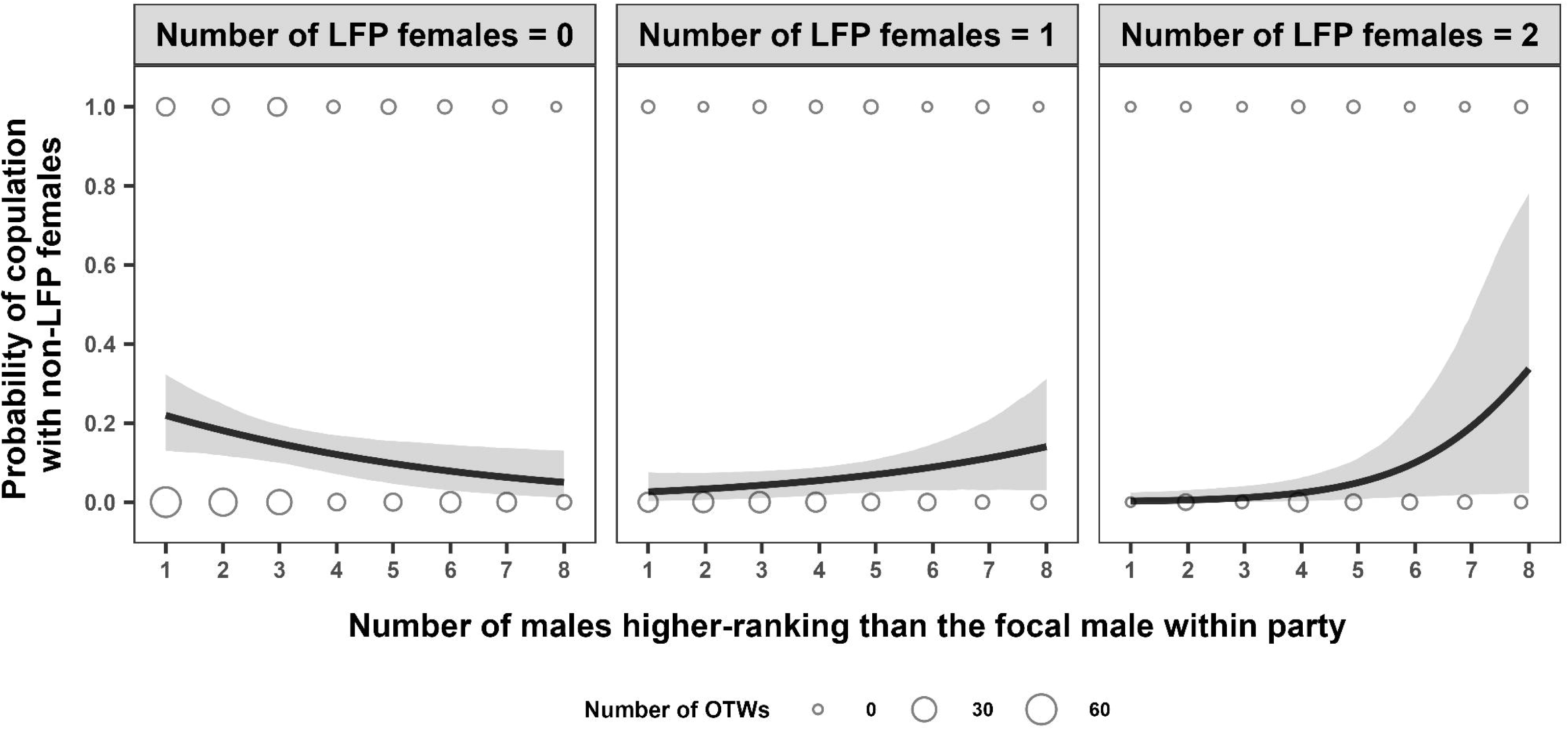
Relationship between within-party male dominance rank and the probability of copulation with females outside the likely fertile period (LFP), changing with the number of LFP females. Predicted copulation probabilities are shown separately for one-hour time windows (OTWs) containing zero (left panel), one (center panel), or two (right panel) females in the LFP. The size of open circles indicates observed total counts of whether copulation occurred for each OTW. Solid lines show model-predicted probabilities, and shaded areas represent 95% confidence intervals on the probability scale.

## 4. Discussion

### 4.1 Rank-independent male copulation frequency with an increased number of receptive females

All-day focal follows of male bonobos revealed no detectable relationship between male dominance rank and copulation frequency with either adult or adolescent females. This finding contrasts with prior reports of higher copulation rates for dominant males (Kano 1996; Hohmann and Fruth 2003; Surbeck, Mundry, and Hohmann 2011; Yokoyama and Furuichi 2022; Ryu et al. 2025). One plausible explanation for this discrepancy is methodological heterogeneity among studies. These previous studies were conducted under provisioning or relied on short-duration focal follows (10–60 minutes). Such approaches can over-represent centrally positioned individuals and larger parties, thereby missing copulations occurring at the periphery or during small-party ranging (Altmann 1974; Drickamer 1974). This concern is particularly relevant if subordinate males pursue mating opportunities while avoiding interference by dominants (Furuichi and Ihobe 1994; Kano 1996; this study). Under these condition, all-day focal follows are more likely to yield less biased estimates of mating activity across ranks. Nevertheless, because our inference is based on six months of observations in a single group, additional multi-group and long-term datasets using comparable all-day protocols are needed to assess the generality of rank-independent copulation frequency.

Across group-living mammals, male dominance rank may predict mating success when ovulation timing is temporally asynchronous among females and the operational sex ratio is male-biased (i.e., when few females are receptive relative to the number of mature males) (Emlen and Oring 1977; Clutton-Brock 1988). Such conditions can generate strong reproductive skew if dominant males exclude subordinates from mating opportunities. In our study, however, multiple receptive females were often simultaneously present (mean = 2.9 females per one-hour time window), a context that likely dispersed male mating effort and reduced the scope for rank-based monopolization. This interpretation is consistent with the expectation that female reproductive synchrony weakens priority-of-access dynamics by increasing the number of simultaneously receptive females (Bulger 1993; Ostner, Nunn, and Schulk 2008; Soulsbury 2011), and with evidence from seasonally breeding mammals in which high synchrony can reduce mating skew (e.g., Japanese macaques, *Macaca fuscata*: Soltis, Thomsen, and Takenaka 2001, rhesus macaques, *Macaca mulatta*: Dubuc et al. 2011; Barbary macaques, *Macaca sylvanus*: Bissonnette, Bischofberger, and van Schaik 2011; Tibetan macaques, *Macaca thibetana*: Zhang et al. 2024, raccoons, *Procyon lotor*: Gehrt and Fritzell 1999; Soay sheep, *Ovis aries*: Preston et al. 2005).

### 4.2 Rank-biased access to females in the likely fertile period and predictability of ovulation from the visual signals

Despite the absence of rank effects on males’ copulation frequencies, high-ranking males accounted for a disproportionate share of copulations with females in the likely fertile period (LFP), based on both female-based focal follows and ad libitum records. This pattern is consistent with genetic evidence for pronounced male reproductive skew in bonobos (Surbeck et al. 2017; Ishizuka et al. 2018; Mouginot, Cheng, and Wilson 2023) and with behavioral and hormonal evidence that males can extract usable information about ovulation timing from graded swelling cues and concentrate mating effort in periovulatory periods (Ryu et al. 2025). Although overt, direct disruption of copulations by dominant males was infrequent in our observations, dominants may nonetheless have constrained subordinates’ access to LFP females through close following and positional control. Comparable patterns have been reported in other non-human primates in which females remain receptive across extended period that includes infertile days, yet male mating effort is biased toward the periovulatory phase (e.g., chimpanzees: Deschner et al. 2004; mandrills, *Mandrillus sphinx*: Setchell, Charpentier, and Wickings 2005; savannah baboons, *Papio cynocephalus*: Alberts, Buchan, and Altmann 2006; sifakas, *Propithecus verreauxi*: Mass, Heistermann, and Kappeler 2009; rhesus macaques: Dubuc et al. 2012; crested macaques, *Macaca nigra*: Higham et al. 2021). Together, these findings underscore that variation in male reproductive success can be shaped more by access to females within the fertile window than by total copulation counts.

In addition to male rank, maternal effects may have reinforced the rank bias in access to LFP females in this group. The alpha male (KT) showed the highest proportion of copulations with LFP females, and his maternal brothers (KY and NB), all sons of the alpha female (Ki), also showed much higher proportions than the across-male median. We also observed Ki supporting her sons during conflicts with other males, including one instance in which she intercepted another male’s copulation attempt. This pattern is consistent with evidence that the presence of mothers, and particularly their dominance status, enhances sons’ mating and reproductive success in bonobos by increasing access to females and enabling direct maternal intervention during male-male competition (Kano 1996; Furuichi 1997; 2011; Surbeck, Mundry, and Hohmann 2011; Surbeck et al. 2019; Yokoyama and Furuichi 2022; Shibata and Furuichi 2024). In contrast, the elevated LFP access of these maternal brothers is unlikely to reflect cooperative alliances among them, because aggressive interactions were most frequent within this set of brothers and coalitionary support was rare in our observations. Taken together, these results suggest that rank-biased access to females in the fertile window may be reinforced by maternal support rather than by coalition formation among maternal brothers.

Because ejaculates are energetically costly to produce (Dewsbury 1982; Thomsen et al. 2006; Parker 2016), males are expected to allocate mating effort strategically by prioritizing copulations with females in the periovulatory period while reducing ejaculatory investment in non-conceptive copulations (e.g., Aujard et al. 1998; Li, Yin, and Zhou 2007; Heistermann et al. 2008; Garcia, Shimizu, and Huffman 2009). In contrast to a strictly deterministic allocation rule, our observations indicate that even the alpha male bonobo copulated with non-LFP females in nearly half of his observed copulation events, and ejaculation was occasionally confirmed during copulations with non-LFP females. These results suggest that male bonobos’ assessment of female fertility is probabilistic rather than categorical. At the same time, our findings are broadly consistent with conditional allocation: within parties, higher-ranking males were less likely to copulate with non-LFP females when LFP females were present, and ejaculation was observed approximately twice as often during copulations with LFP females as during copulations with non-LFP females, although this difference was not statistically significant. Taken together, these patterns suggest that male bonobos bias mating effort toward females estimated to have higher conception probability while maintaining occasional ejaculatory investment in females with lower conception probability, plausibly reflecting uncertainty in fertility assessment rather than indiscriminate mating.

In this study, female fertility status was inferred from daily swelling scores by operationally defining the LFP based on the empirical relationship between detumescence timing and hormone-estimated ovulation reported by Ryu et al. (2025). This operational definition yielded a rank-biased pattern of copulations with LFP females that is congruent with paternity pattens reported by genetic studies of bonobos. Thus, the swelling-based proxy can be useful for estimating the fertile window when intensive hormonal sampling is infeasible. However, because swelling patterns can vary substantially in their relationship to ovulation in bonobos, the LFP should be interpreted as an approximate indicator rather than a precise fertile window.

### 4.3 Within-party rank dynamics and alternative mating strategies by subordinate males

Party-level analyses provide additional insight into how male bonobos allocate mating effort on short temporal scales. Although within-party male dominance rank did not affect the probability of copulation per se, it strongly influenced the fertility status of mating partners. Higher-ranking males preferentially copulated with LFP females, whereas lower-ranking males increasingly copulated with non-LFP females as the number of LFP females in the party increased. This pattern suggests that subordinate males can secure mating opportunities by targeting females that are less likely to be guarded by dominant males. Such an alternative tactic may increase subordinates’ participation in sperm competition, even if it rarely translates into paternity. Furthermore, over longer time horizons, avoiding direct contests with dominant males may also reduce injury risk and extend reproductive lifespan. This could be especially relevant in bonobos, where female dominance rank tends to increase with age and maternal rank can shape sons’ future competitive prospects (Furuichi 1997; Tokuyama and Furuichi 2016; Toda and Furuichi 2020; Shibata and Furuichi 2023).

Comparable patterns have been reported in chimpanzees, where subordinate males maintain mating activity by shifting effort toward females that are less closely guarded by dominant males (Nishida 1997; Matsumoto-Oda 1999; Deschner et al. 2004). Unlike bonobos, however, subordinate chimpanzee males sire a non-trivial proportion of offspring (Constable et al. 2001; Boesch et al. 2006; Wroblewski et al. 2009; Newton-Fisher et al. 2010). One tactic subordinate males adopt is to form a temporary exclusive association with a given female by leading her away from other males and to mate with her repeatedly over several days during the fertile phase (i.e., consortship; Tutin 1979; Matsumoto-Oda 1999; Constable et al. 2001; Wroblewski et al. 2009; Newton-Fisher et al. 2010; Bray, Pusey, and Gilby 2016). The fission-fusion grouping pattern may afford subordinate males more opportunities to practice the consortship tactic, particularly in chimpanzee groups where females spend more time alone (e.g., Gombe), than in more spatially cohesive groups such as baboons and macaques (Wroblewski et al. 2009). However, consortship does not appear to play a major role in bonobo reproduction (Kano 1992; Takahata, Ihobe, and Idani 1999; Surbeck et al. 2017). In bonobos, the presence of maximally tumescent females serves as a key driver increasing female-female gregariousness (Surbeck et al. 2021). This social context may make it easier for males to locate and monitor receptive females, potentially increasing the extent to which dominant males can monopolize reproductive opportunities (Mouginot, Cheng, and Wilson 2023).

Even if the ultimate reproductive payoff of alternative tactics differs between species, subordinate males in bonobos may have more frequent opportunities to obtain mating partners than subordinate males in chimpanzees. because bonobo parties often include more receptive females relative to the number of mature males (Furuichi and Hashimoto 2002; Surbeck et al. 2021). In our study, 92.6% of all-day focal follows of male bonobos (38 of 41 days) in the E1 group included at least one maximally tumescent female, which was more than twice the percentage reported for all-day focal follows of male chimpanzees in the Kanyawara group (38.8%; 33 of 85 days; Georgiev et al. 2014). In addition, the mean number of maximally tumescent females present on a given day was substantially higher in E1 bonobos (4.1 ± 2.4 SD) than Kanyawara chimpanzees (0.5 ± 0.1 SE). Although the E1 group during our study period contained the largest number of females ever (Figure S1), these comparisons suggest that the earlier resumption of sexual receptivity postpartum in female bonobos than chimpanzees may provide mature males of all ranks with frequent mating opportunities.

### 4.4 Implications for female mate choice and potential benefits from prolonged sexual signaling

Together, our results indicate that pronounced reproductive skew among male bonobos is shaped primarily through rank-biased access to fertile females, potentially reinforced by maternal support and high female gregariousness. Female mate choice may also contribute to this skew in bonobos, where females can ignore or refuse male courtship (Furuichi and Hashimoto 2004; Hashimoto and Furuichi 2006) and sexual coercion appears to be substantially limited (Paoli 2009). If male dominance rank covaries with aspects of phenotypic quality, such as disease resistance, weapon traits, or behavioral dispositions, female preferences for dominant males could yield indirect fitness benefits consistent with good genes models (Kirkpatrick 1996; Møller and Alatalo 1999). Comparative evidence from other primates suggests that male rank can correlate with traits plausibly linked to health condition or competitive ability, including immune function and oxidative status (rhesus macaques: Georgiev et al. 2015), canine dimensions (anubis baboons, *Papio anubis*: Galbany et al. 2015), and personality profiles (chimpanzees: Weiss et al. 2023). Identifying whether dominance rank in male bonobos tracks measurable indicators phenotypic quality will be important for evaluating the role of female mate choice in generating male reproductive inequality.

Our findings also support the updated prolonged sexual receptivity hypothesis that a higher number of receptive females relative to the number of mature males disperses male mating effort across multiple females and thereby mitigates male-male contest competition over mates. Females may benefit from weakening male-male contest competition, potentially reducing risks linked to infanticide and/or sexual coercion. However, prolonged sexual receptivity in female bonobos likely has functions beyond shaping male mating strategies. Maximally tumescent females attract other females as well as males, increasing female-female gregariousness (Surbeck et al. 2021) and eliciting affiliative interactions among females (Ryu, Hill, and Furuichi 2015; Anzà, Demuru, and Palagi 2021). Moreover, female bonobos frequently form coalitions during aggressive interactions and maintain high dominance relative to males (Tokuyama and Furuichi 2016; Samuni and Surbeck 2023; Surbeck et al. 2025). Given that strong social bonds among unrelated females is a central feature of bonobo society (Moscovice et al. 2017; Furuichi 2023; Samuni and Surbeck 2023), prolonged sexual receptivity may have been favored primarily because it promotes female-female affiliation and cooperation, with any reduction in male-male contest competition emerging as a secondary consequence.

This study adds to evidence that female bonobos remain receptive for prolonged periods, yet ovulation cues are sufficiently informative for males to bias mating effort toward the fertile phase (Ryu et al. 2025) and for paternity to be strongly skewed toward dominant males (Mouginot et al. 2024). This combination is not necessarily incompatible with a role of prolonged sexual receptivity in paternity confusion. Even when paternity is concentrated among dominant males, frequent non-conceptive copulations with multiple subordinates may still create uncertainty about paternity and reduce incentives for aggression toward infants (Palombit 2015). In parallel, prolonged sexual receptivity can yield additional benefits by facilitating socio-sexual interactions that strengthen affiliation and cooperation among unrelated females (Ryu, Hill, and Furuichi 2015; Moscovice et al. 2019; Surbeck et al. 2021; Furuichi 2023). An unresolved question is whether females obtain additional benefits from males through non-conceptive copulations, such as increased tolerance around food resources. Although anecdotal, bonobos have been reported to exchange sex for food (Kuroda 1984; Van Krunkelsven et al. 1996; but Yamamoto 2015). Further evidence on the benefits females receive through non-conceptive sexual interactions may help explain interspecific variation in the duration and dynamics of sexual receptivity among anthropoid primates.

## 5. Acknowledgements

We thank the staff of the Centre de Recherche en Écologie et Foresterie (CREF), Democratic Republic of the Congo, for supporting our fieldwork at Wamba in the Luo Scientific Reserve—especially Messrs. Bahanande Iyokango, Yembo Bafaluka, Batsina Bambambe, and Bakka Batuafe. We are also grateful to the local assistants who helped with our fieldwork and to the residents of Wamba for their hospitality. Fieldwork by K. Toda was supported by Grant-in-Aid for Scientific Research (21K13670) from the Japan Society for the Promotion of Science and by the Canon Foundation in Europe–Kyoto University Japan–Africa Exchange Fellowship.

## Ethics statement

Our field studies of free-ranging bonobos at Wamba in the Luo Scientific Reserve were approved by the Ministry of Scientific Research and the Center for Research on Ecology and Forestry in the Democratic Republic of Congo. This study adhered to the Animal Research Guidelines of The Graduate University for Advanced Studies (SOKENDAI), Japan, revised on March 23, 2021.

## Data availability statement

The data that support the findings of this study are available from the corresponding author upon reasonable request.

