## Supplementary Figure S1-S8 for "Rank-biased access to females in the likely fertile period despite comparable copulation rates in male bonobos at Wamba": 5th_supplementary_260209.docx

**Supporting Information**

**FIGURE S1** Demographic changes in the E1 group at Wamba between 1976 and 2023. It was not available about group compositions of this group between 1997 and 2003 due to the lack of observations.

**FIGURE S2** Diagnostics for scaled residuals by simulating from the Poisson GLMM testing the effect of male rank on counts of copulations with adult females

**FIGURE S3** Diagnostics for scaled residuals by simulating from the Poisson GLMM testing the effect of male rank on counts of copulations with adolescent females

**FIGURE S4** Diagnostics for scaled residuals by simulating from the binomial GLMM examining the effect of female LFP status on the proportion of copulations with high- versus low-ranking males

**FIGURE S5** Diagnostics for scaled residuals by simulating from the binomial GLM examining the effect of male rank on the proportion of copulations with LFP versus non-LFP females

**FIGURE S6** Diagnostics for scaled residuals by simulating from the binomial GLMM examining the effect of the number of males higher ranking than the focal male on the occurrence of copulations within (OTWs)

**FIGURE S7** Diagnostics for scaled residuals by simulating from the binomial GLMM examining the effect of within-party male rank on the occurrence of copulations with LFP females within OTWs

**FIGURE S8** Diagnostics for scaled residuals by simulating from the binomial GLMM examining the interaction between within-party male rank and the number of LFP females present on the occurrence of copulations with non-LFP females within OTWs
