## Supplementary figures and images for "Rank-biased access to females in the likely fertile period despite comparable copulation rates in male bonobos at Wamba"

### FIGURE S1.png

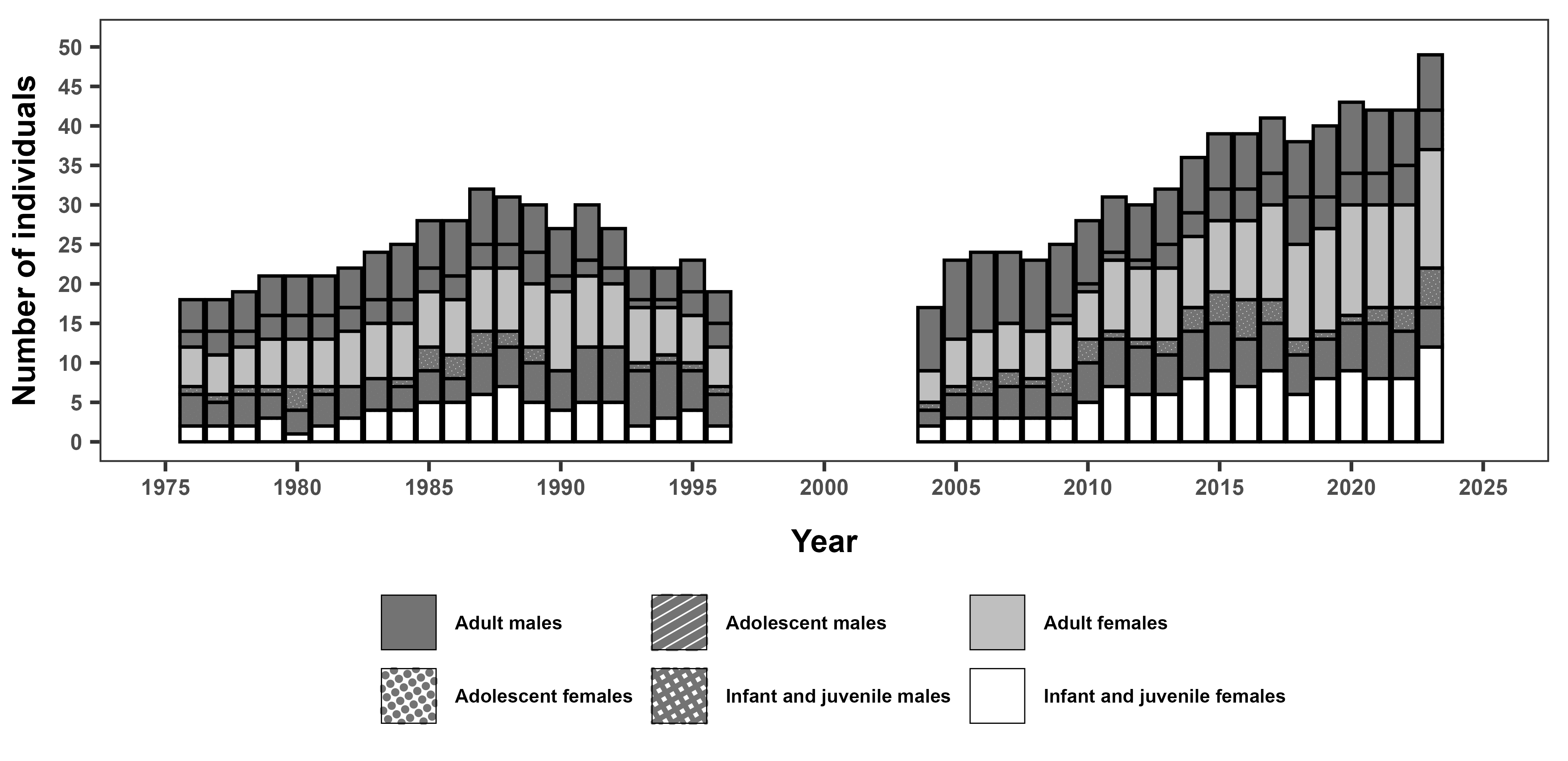

### FIGURE S2.png

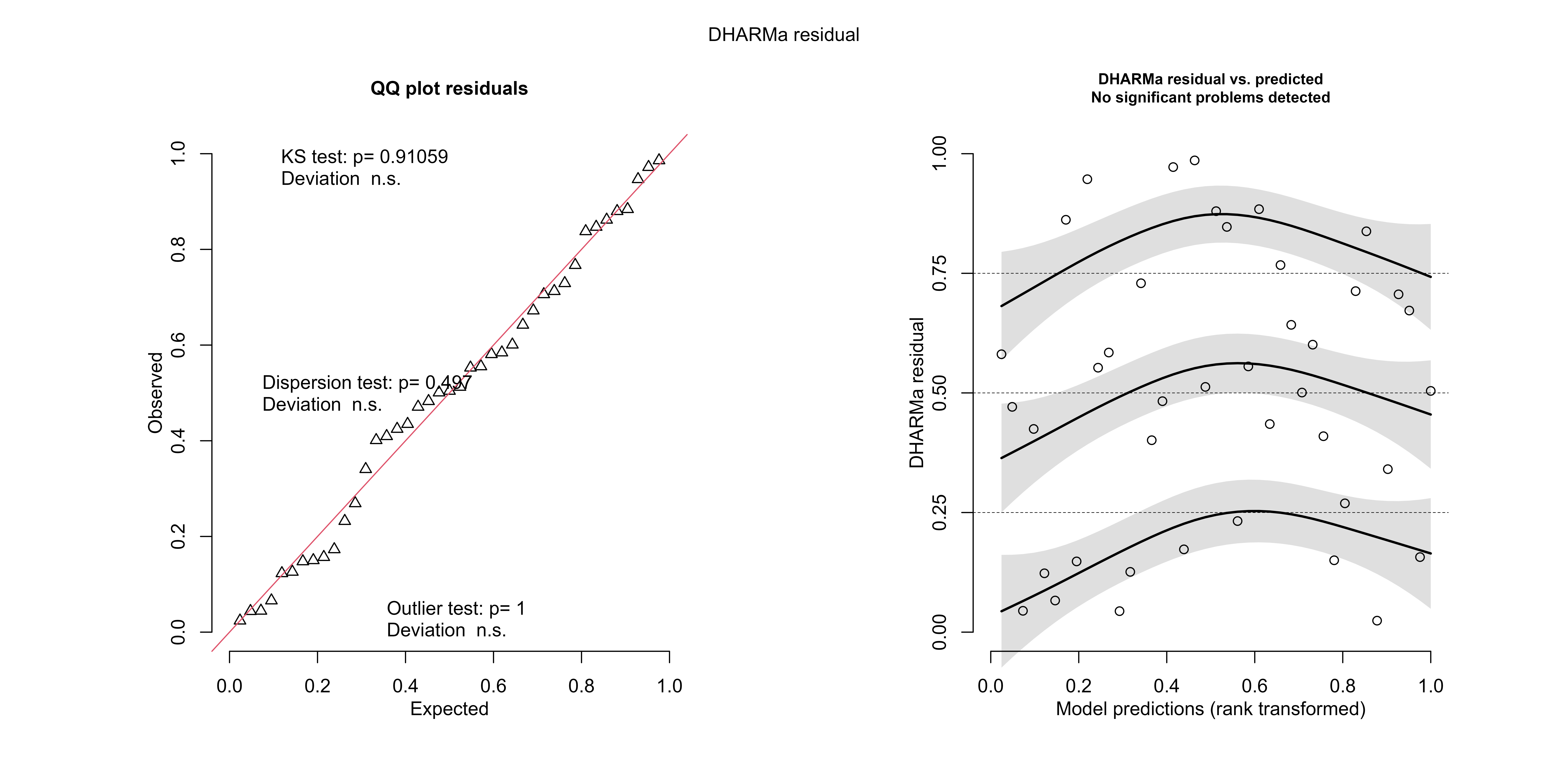

### FIGURE S3.png

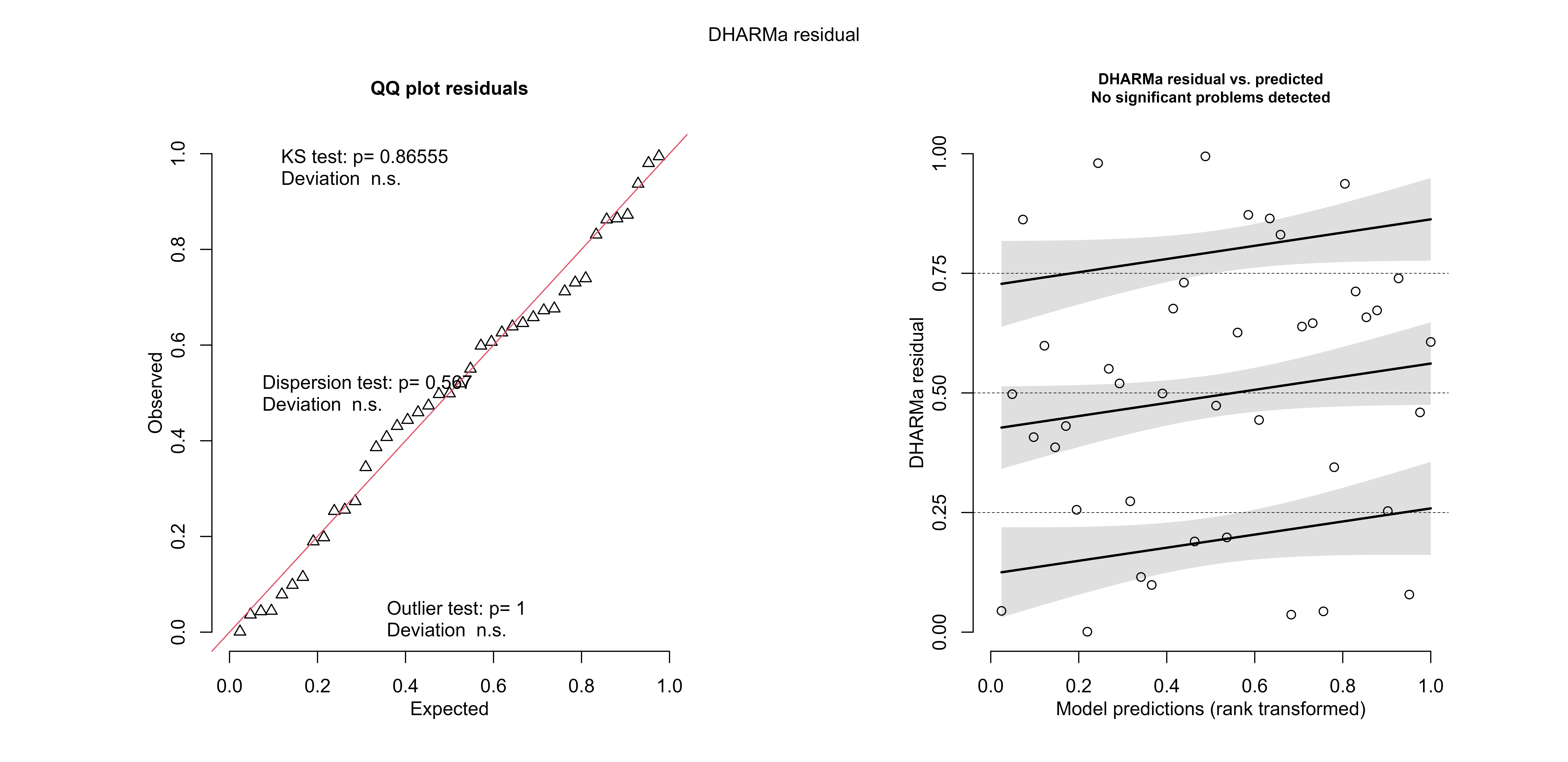

### FIGURE S4.png

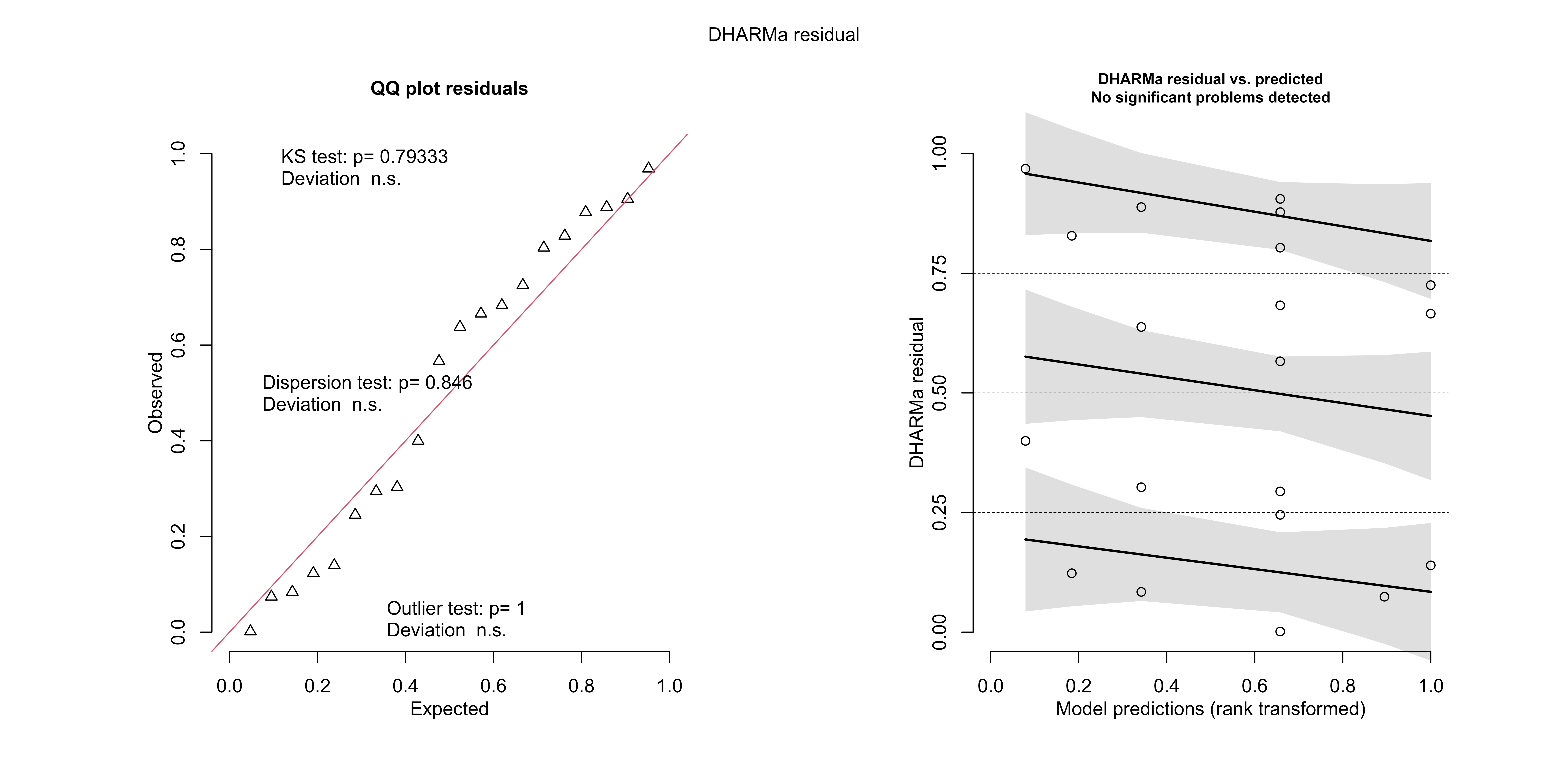

### FIGURE S5.png

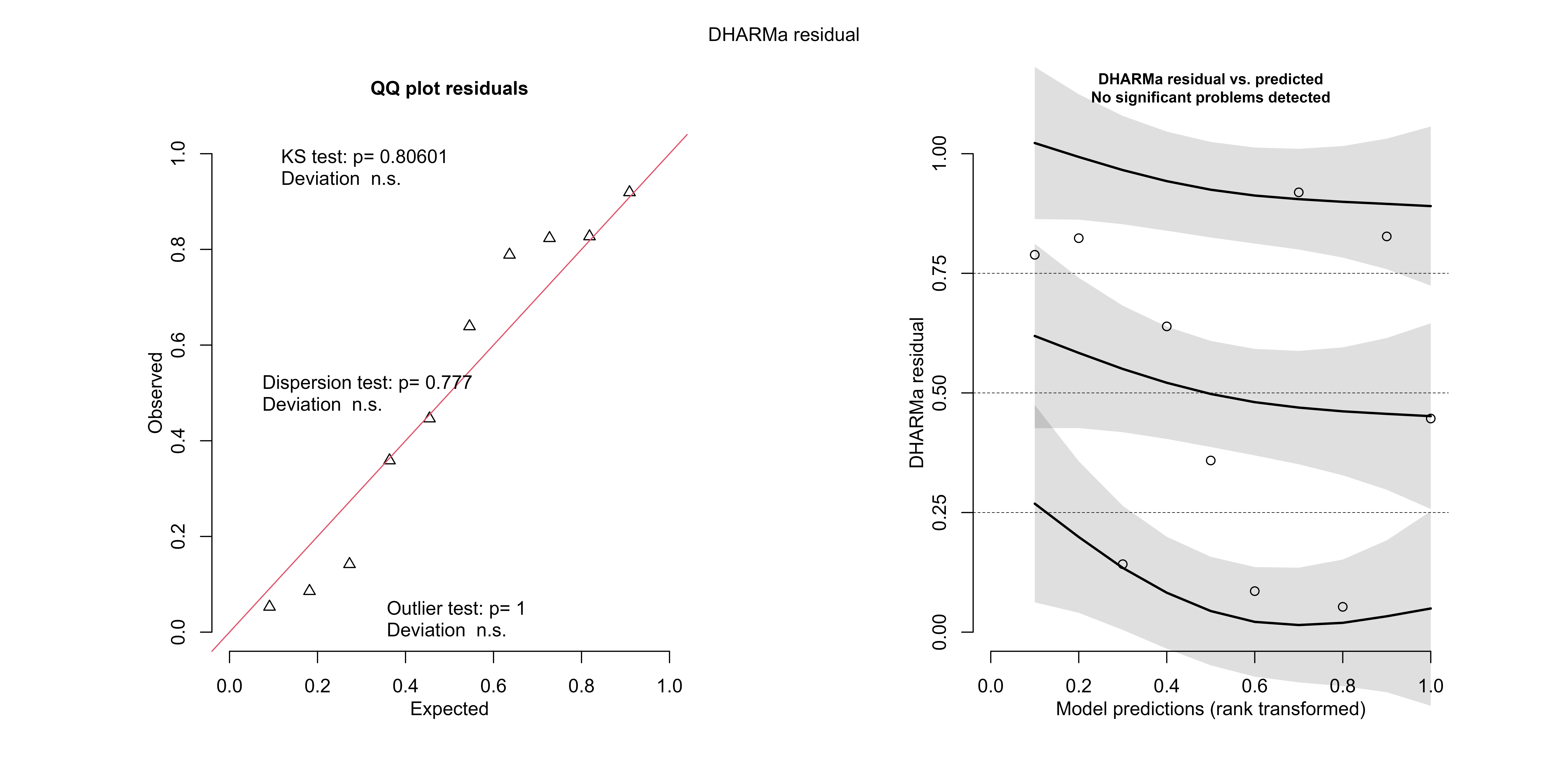

### FIGURE S6.png

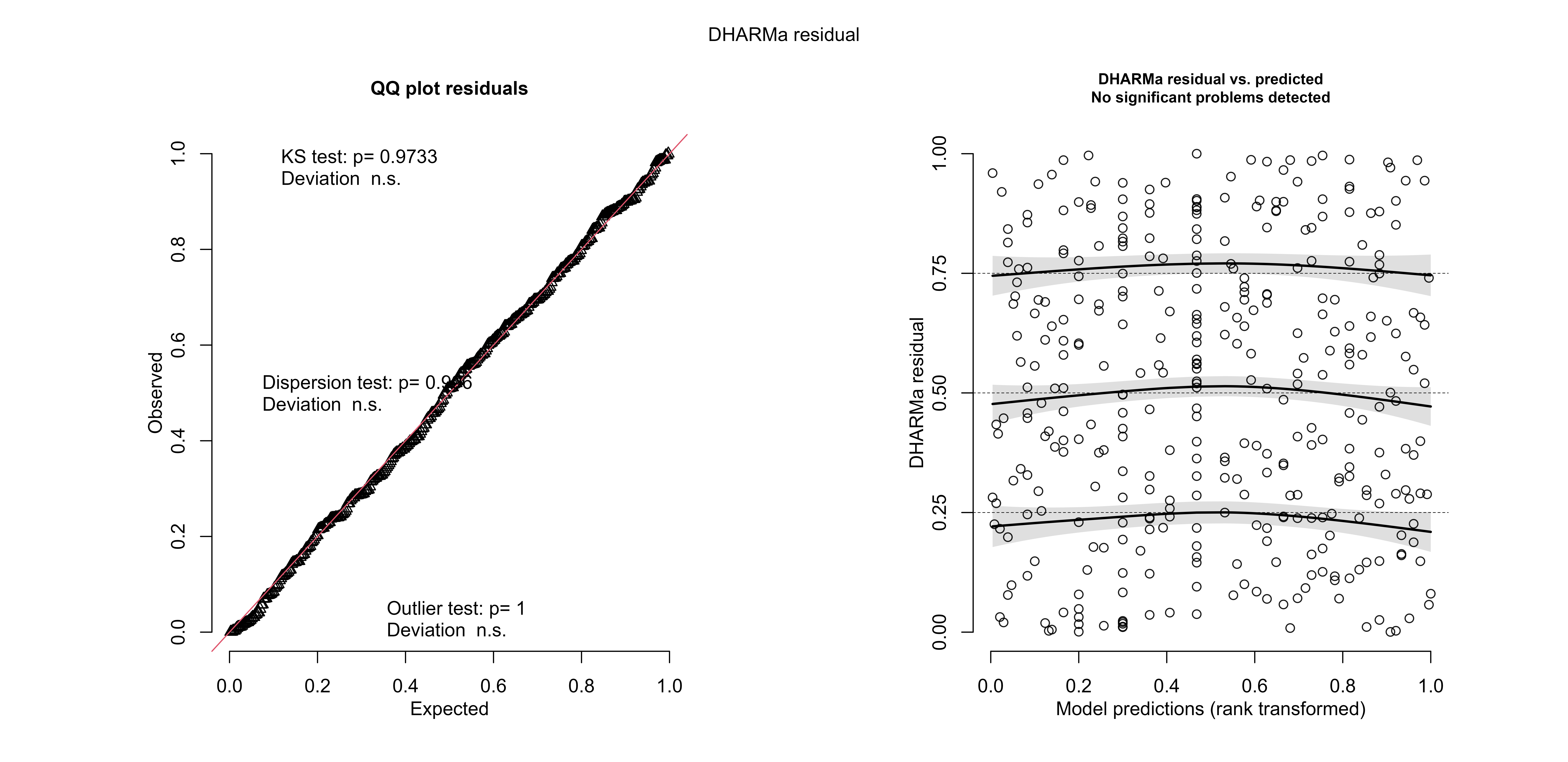

### FIGURE S7.png

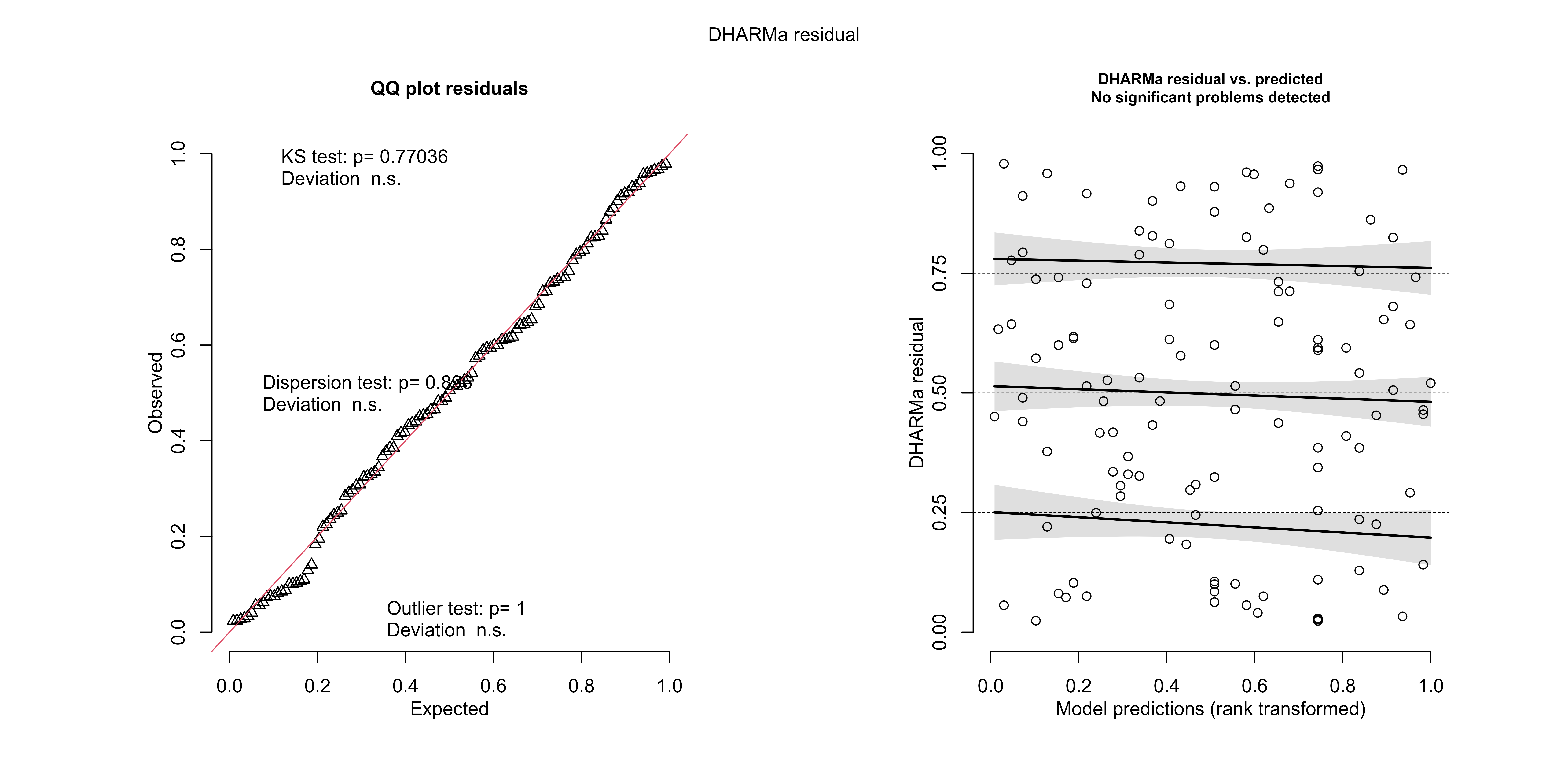

### FIGURE S8.png

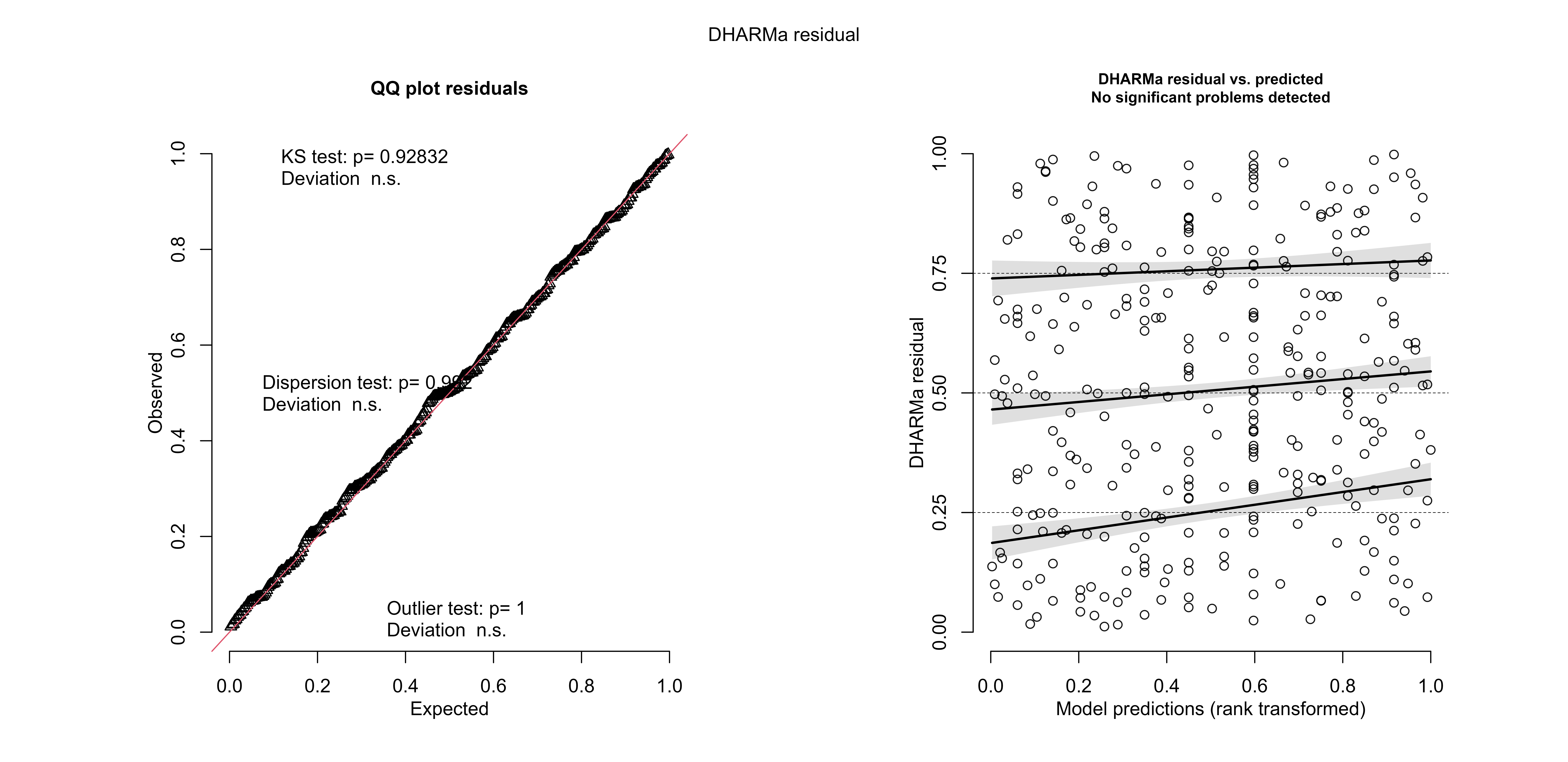
